# Blood flow coordinates collective endothelial cell migration during vascular plexus formation and promotes angiogenic sprout regression via *vegfr3/flt4*

**DOI:** 10.1101/2021.07.23.453496

**Authors:** Yan Chen, Aaron M. Savage, Zhen Jiang, Hyejeong R. Kim, Paul C. Evans, Robert N. Wilkinson

## Abstract

Nascent vascular networks adapt to the increasing metabolic demands of growing tissues by expanding via angiogenesis. As vascular networks expand, blood vessels remodel, progressively refining vascular connectivity to generate a more haemodynamically efficient network. This process is driven by interplay between endothelial cell (EC) signalling and blood flow. While much is known about angiogenesis, considerably less is understood of the mechanisms underlying vessel remodelling by blood flow. Here we employ the zebrafish sub-intestinal venous plexus (SIVP) to characterise the mechanisms underlying blood flow-dependent remodelling. Using live imaging to track ECs we show that blood flow controls SIVP remodelling by coordinating collective migration of ECs within the developing plexus. Blood flow opposes continuous ventral EC migration within the SIVP and is required for regression of angiogenic sprouts to support plexus growth. Sprout regression occurs by coordinated polarisation and migration of ECs from non-perfused leading sprouts, which migrate in opposition to blood flow and incorporate into the SIV. Sprout regression is compatible with low blood flow and is dependent upon *vegfr3/flt4* function under these conditions. Collectively, these studies reveal how blood flow sculpts a developing vascular plexus by coordinating EC migration and balancing vascular remodelling via *vegfr3/flt4*.

## Introduction

During early development, cellular uptake of nutrients and elimination of metabolic waste must occur prior to the formation of specialised organs responsible for these functions. Consequently, the nascent vascular networks that mediate metabolic exchange must undergo rapid adaptation to accommodate the dynamic metabolic demands of growing tissues. As vascular networks expand, they must optimise metabolic exchange while minimising resistance to blood flow. Evolution has therefore favoured mechanisms that enhance haemodynamic efficiency (Campinho et al., 2020). Angiogenic sprouting must be balanced with vascular remodelling to maintain haemodynamic efficiency during network expansion. Remodelling occurs either by fusion or regression of angiogenic sprouts to form functional tubular vessels, or the pruning of redundant blood vessels (Ribatti and Crivellato, 2012). Endothelial cells (ECs), which line the inner surface of blood vessels play a central role in this process by responding dynamically to changes in their local environment (Udan et al., 2013). While most blood vessels develop from pre-existing patent vessels with stable blood flow, our understanding of the mechanisms governing vascular remodelling and the influence of blood flow on these processes remains incomplete. For example, it is not yet clear whether vascular regression is driven by active signalling pathways, the withdrawal of survival cues, or a combination of both, nor whether these mechanisms vary across vascular beds (Korn and Augustin, 2015).

The vascular network of the zebrafish sub-intestinal venous plexus (SIVP) forms bilaterally in the embryo and is structurally similar to the mammalian vitelline veins which connect the embryo with extraembryonic circulation in the yolk sac (Goi and Childs, 2016). Owing to its developmental plasticity the SIVP has become a valuable model for investigating the cellular mechanisms underlying vascular remodelling (Goi and Childs, 2016; Hen et al., 2015; Koenig et al., 2016; Lenard et al., 2015). The SIVP initially facilitates nutrient uptake from the yolk and later contributes to vascularisation of the larval the digestive system (Isogai et al., 2001). The SIVP comprises the supra-intestinal artery (SIA), bilateral sub-intestinal veins (SIV) and inter-connecting vessels (ICVs) which connect the SIA and SIV (Fig. 1A). The SIVP originates from angioblasts in the ventral posterior cardinal vein, which give rise to both the SIA and the SIV (Goi and Childs, 2016; Hen et al., 2015; Koenig et al., 2016). The primitive SIVP expands bilaterally over the yolk surface and ECs from the primary SIV migrate dorsally throughout the plexus to form branches and the SIA (Goi and Childs, 2016; Hen et al., 2015; Koenig et al., 2016). In mammals, progressive refinement of a vascular network by vessel pruning or regression involves dynamic migration and rearrangement of ECs (Franco et al., 2016, 2015; Udan et al., 2013). ECs are capable of migrating within patent blood vessels during vascular network formation (Christ et al., 1990; Franco et al., 2015) leading us to hypothesise that coordinated EC motility is essential for SIVP remodelling and may be regulated by blood flow.

**Figure 1.**
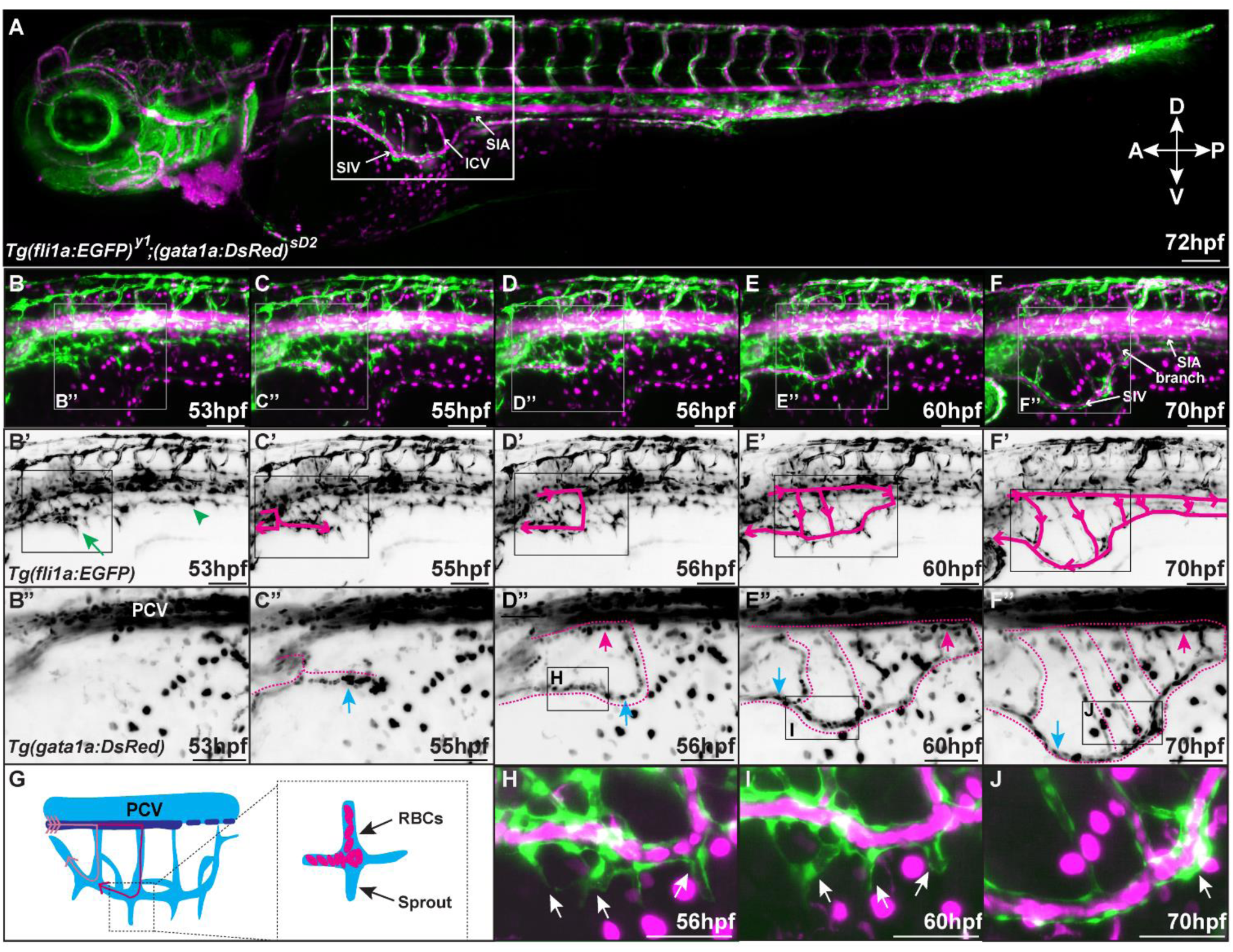
SIVP perfusion is coincident with leading sprout regression. (**A-F**) Representative images taken from a time-lapse (Supplementary Movie 1) between 53-70hpf showing perfusion of the sub-intestinal venous plexus (SIVP). *Tg(fli1a:EGFP)^y1^*was used to label endothelial cells (green) and *Tg(gata1a:DsRed)^sD2^*labels erythrocytes (magenta). Region highlighted in A is displayed in B-F, and region highlighted in B-F is enlarged in B’-F’. Region highlighted in B’-F’ is enlarged in B’’-F’’ and displays the path of circulating erythrocytes. Circulation enters the anterior SIVP between 53hpf (**B-B”**) and 55hpf (**C-C’’**) and perfuses the SIV from anterior to posterior as it develops (**D-F’’**). Blood enters the SIVP from the supraintestinal artery (SIA) (**D’-F’’**, magenta arrows) an extension of the upstream anterior mesenteric artery, through the SIVP branches, drains into the SIV (**C’’-F’’**, blue arrows) and exits the SIV via the hepatic portal vein (HPV). **G**) Schematic representation of SIVP perfusion. **H-J**) Leading sprouts are observed to regress and undergo anastomosis in the presence of blood flow (arrows). ICV, inter-connecting vessel; PCV, posterior cardinal vein; RBC, red blood cells; SIA, supra-intestinal artery; SIV, sub-intestinal vein. Scale bars = 100µm (A-F’), 50µm (B’’-J).

Vascular Endothelial Growth Factor (VEGF) signalling controls diverse aspects of EC biology, including migration and survival (Akeson et al., 2010; Ferrara et al., 2003). VEGF ligands signal via their cognate receptor tyrosine kinase receptors VEGFR1-3, with VEGFR2 serving as the principal signalling receptor in mammalian blood ECs (Simons et al., 2016). In zebrafish, Vegfr4/Kdrl and Vegfr2/Kdr are functionally analogous to mammalian VEGFR2 (Bussmann et al., 2008). ECs sense blood flow via junctional mechanosensory complexes, most notably the VE-cadherin, PECAM-1, VEGFR2/3 complex, which operates independently of VEGF ligands and is essential for arterial vascular remodelling (Coon et al., 2015; Tzima et al., 2005). Endothelial VEGFR3/Flt4 levels establish a fluid shear stress set point, and sustained deviations from this threshold trigger vascular remodelling in arteries (Baeyens et al., 2015; Tzima et al., 2005). Zebrafish *vegfaa* mutants exhibit severe defects in SIVP formation (Habeck et al., 2002; Koenig et al., 2016) and combined loss of *kdr* and *kdrl* similarly disrupts SIVP development (Habeck et al., 2002; Koenig et al., 2016). Sprout pathfinding within the SIVP depends on the guidance receptor, PlexinD1, and plexus expansion is constrained by inhibition of Bone Morphogenetic Protein (BMP) signalling (Goi and Childs, 2016). SIVP remodelling occurs via flow-dependent pruning and fusion of vessel branches (Hen et al., 2015; Lenard et al., 2015). Retraction of SIVP leading sprouts has been observed (Hen et al., 2015), but whether this process is flow-dependent remains unknown. Moreover, a systematic analysis of EC migration during SIVP formation is lacking and the interplay between blood flow and signalling pathways in co-ordinating plexus remodelling remains poorly understood.

Here we demonstrate that blood flow plays a critical role in coordinating EC migration during vascular plexus formation and remodelling. Through time-lapse imaging and quantitative analysis, we show that blood flow orchestrates collective EC migration throughout the developing SIVP and promotes regression of leading angiogenic sprouts to support plexus expansion. Our data suggest that sprout regression occurs by coordinated EC rearrangement between non-perfused sprouts and the vessel lumen, driven by directed EC polarisation and migration against blood flow. Using erythrocyte depletion to reduce circulation, we find that sprout regression can occur under low flow conditions, but this process requires *flt4* function. Together, these findings provide a clear view of the complex cellular dynamics underlying vessel remodelling during vascular plexus formation and highlight the central role of EC responses to blood flow.

## Methods

### Zebrafish strains

Maintenance of zebrafish and experimental procedures involving zebrafish were carried out according to UK national guidelines and under UK Home Office licenses. The following zebrafish lines were employed in this study: *Tg(fli1a:*EGFP*)^y1^*(Lawson and Weinstein, 2002)*, Tg(gata1a:*DsRed*)^sD2^* (Traver et al., 2003), *Tg(fli1a:AC-TagRFP)^SH511^*(Savage et al., 2019), *Tg(kdrl:EGFP)^s843^* (Jin et al., 2005), *Tg(fli1a:nls-*mCherry*)^SH550^*(this study), *Tg(fli1a:nls-*EGFP*)^SH549^* (this study), *Tg(fli1a:golgi-*TagRFP*; cryaa:*CFP*)^SH529^* (this study), *flt1*^bns29^ (Matsuoka et al., 2016), *flt4^qmc316^*, (this study).

### Generation of Transgenic lines

*Tg(fli1a:nls-*EGFP*)^SH549^* and *Tg(fli1a:nls-mCherry)^SH550^* were generated using the Tol2 Kit (Kwan et al., 2007) via standard methods as previously described (Savage et al., 2019) using the following components: p5E-fli1aep (Villefranc et al., 2007) pME-mCherry or pME-EGFP, p3E-SV40pA, and pDestTol2-pA2. *Tg(fli1a:golgi-TagRFP, cryaa:cerulean)^SH529^* was generated by fusing amino acids 1-60 of Human B4GALT1 to the N-terminus of TagRFP via *in silico* synthesis. attB1/attB2R sites were added to the fusion product by PCR (Supplementary Table 1) to generate pME-Golgi-TagRFP. The *fli1a:golgi*-TagRFP*; cryaa:*CFP construct was generated using the Tol2 Kit and the following components: p5E-fli1aep, pME-Golgi-TagRFP, p3E-SV40pA, and pDestTol2cryCFP. Embryos were injected at one-cell stage with 25 ng/μl *Tol2* mRNA and corresponding plasmid DNA.

### Microinjection of morpholinos and mRNA

Microinjections were performed on single cell embryos with 1nl injection volume. Embryos were injected with 0.8ng morpholinos (Supplementary Table 1) including control, *tnnt2a* (Sehnert et al., 2002), *flt4* (Hogan et al., 2009b), and *gata1a* (Galloway et al., 2005) or 200pg mRNA including *mTurquoise2* and *vegfaa_165_* (Lawson et al., 2002).

### G0 *tnnt2a* CRISPR mutants

Target sequences were ordered as crRNAs (Merck) based on previously published sequences (Wu et al., 2018) (Supplementary Table 1) and co-injected alongside tracrRNA (Merck) in equimolar ratios and EnGen®Spy Cas9 NLS protein (NEB) into 1-cell zebrafish embryos.

### RNA *in situ* hybridisation

Alkaline phosphatase wholemount *in situ* hybridisation experiments were performed using standard methods as described previously (Wilkinson et al., 2012) using probes against *mflt1* (Krueger et al., 2011), *sflt1* (Krueger et al., 2011), *flt4* (Thompson et al., 1998), *kdrl* (Fouquet et al., 1997). Detailed protocols are available upon request.

### Live imaging of zebrafish embryos and larvae

Zebrafish were anaesthetised using Tricaine (MS-222, Sigma-Aldrich) and embedded in 1% low-melting point agarose which was held in place in a glass capillary and imaged using a light sheet Z.1 microscope (Zeiss), or Cell Discoverer 7 microscope (Zeiss). The chamber contained E3 and tricaine (164mg/L) to ensure embryos and larvae remained anaesthetised throughout imaging. Imaging was performed at 28.5°C. Images were acquired using ZEN software (Zeiss). Zebrafish were imaged at 5-minute intervals to detect blood flow, and 10-minute intervals to track EC nuclei. To image individual erythrocytes in *gata1a* morphants, time lapses were taken at 50 frames per second for 30 seconds with 1 µm z-stacks. The number of erythrocytes which passed a defined region of the SIV within a 30 second period were manually counted.

### Quantification of SIVP morphology

Image quantification and analysis was performed using FIJI (Schindelin et al., 2012). Parameters employed to quantify SIVP morphology including area, length, vascular loops, sprout number and EC number, were recorded in a region of 5 somite widths. The length of the SIVP was measured as the vertical distance from the SIA to the most ventral part of the SIV. To ensure only endothelial nuclei within the SIVP were quantified, dual channel fluorescence was employed to highlight EC cytosol and nuclei. To quantify average SIV diameter, the SIV was measured at three equidistant points positioned anteroposteriorly and mean value was calculated.

### EC rearrangement and cell trajectory analysis

*Tg(fli1a:nls-*mCherry*)^SH550^* heterozygous embryos were used for cell tracking studies. Time-lapse images were pre-processed using Linear Stack Alignment with SIFT (Lowe, 2004) in FIJI and the movements of EC nuclei were tracked using TrackMate (Tinevez et al., 2017). Any misconnected tracks were manually corrected. Tracking data was exported and analysed using customised MATLAB scripts freely available at https://github.com/yanc0913/SIVP_cell_tracking (Chen et al., 2022). ECs were colour coded as tip cells (magenta), SIVP branches (orange), or SIV cells (blue) depending upon their initial position at the start of tracking (56 hours post fertilisation, hpf). At the end of observation (72 hpf), cell positions were evaluated to determine whether ECs moved between EC subsets e.g., an EC initially located in the SIV which migrated to a branch by the end of tracking was recorded as a single rearrangement event. The number of ECs at initial and final position and the events of EC rearrangement were manually recorded from each time lapse. To analyse migration trajectory, a migration step was defined as one cell at position 1 (x_1_ and y_1_) migrating to its subsequent position 2 (x_2_ and y_2_) in a time range (from t_1_ to t_2_). The distance (or displacement) of the step (or the track) was calculated as. Track distance was the sum of step displacements. Migration velocity was calculated as step displacement divided by time interval, and the mean velocity at a particular time range was the average of each velocity in that range. The angle of each migration step was calculated using the four-quadrant inverse tangent (*atan2)* function in MATLAB, which computed the arctangent of Δy and Δx.

### Quantification of EC Golgi polarity

In sprout cells, the position of the Golgi at either dorsal (45-135°) or ventral (225-315°) end of the elongated EC nucleus was considered polarised. In SIV cells, the position of the Golgi at either anterior (or downstream of blood flow, 135-180°) or posterior (or upstream of blood flow, 0-45°) end of the elongated EC nucleus was considered polarised.

### Generation of *flt4^qmc316^* allele

gRNAs targeting *flt4* (ENSDARG00000104453) were ordered from IDT DNA Technologies and targeted the following sequence 5’-CCCAGGGTATAGAGAAATCAGGC-3’ in exon 3. crRNAs were annealed with tracrRNA (IDT) as previously described (Kroll et al., 2021). Annealed sgRNAs were injected into 1-cell stage zebrafish embryos alongside EnGen® Spy Cas9 NLS (NEB #M0646) in a non-complexed solution. F1 progeny of G0 embryos were genotyped using high resolution melt analysis and PCR using the following primers 5’-AGTGGAGTTTACTGATCGCCA-3’, 5’-CCGTGGTGCCATCAATAACA-3’ to confirm the presence of the *flt4^qmc316^* allele. *flt4^qmc31^* consists of a complex insertion and deletion resulting in a net 132bp insertion, which is predicted to shift frame after 83 residues and truncate following 11 subsequent incorrect residues within the first extracellular Ig-like domain and earlier than previously published *flt4* alleles including a reported null allele within exon 4 (Shin et al., 2016).

### Statistical analyses

Statistical analyses and graph plots were performed using GraphPad Prism. All statistical analysis is described in figure legends, including paired/unpaired *t*-test, ordinary one-way ANOVA, and two-way ANOVA. All error bars display the mean and standard deviation in the figures. *p*-values, unless exact value is listed, are as follows: ns= not significant, *=<0.05, **=*≤*0.01, ***=*≤*0.001, ****=*≤*0.0001.

## Results

### Blood flow controls SIVP remodelling by influencing EC distribution and promoting leading sprout regression

The SIVP develops from angiogenic sprouts which emerge from the posterior cardinal vein (PCV) by 30 hpf and migrate in a ventrolateral direction, bilaterally, over the surface of the yolk (Goi and Childs, 2016; Hen et al., 2015; Koenig et al., 2016). To determine the timing of SIVP development in relation to blood flow, we imaged the developing plexus in the presence of fluorescently labelled erythrocytes (Fig.1A). Between 53 hpf and 55 hpf, blood flow entered the nascent SIVP anteriorly via the supra-intestinal artery (SIA) (Fig. 1A-C”). Circulation progressed from anterior to posterior, and by 72 hpf the plexus became fully perfused (Fig. 1D-F”, G supplementary movie 1). As the SIVP becomes perfused, it undergoes extensive remodelling. Migrating angiogenic sprouts, hereafter referred to as leading sprouts, protrude from the ventral sub intestinal vein (SIV) and were observed to either fuse with neighbouring sprouts or regress, with their ECs reintegrating into the SIV by 70 hpf (Fig. 1H-J, arrows, supplementary movie 1 and 2, arrowheads).

Remodelling of leading sprouts coincided with the onset of blood flow in the SIV. After flow began, new leading sprouts were rarely formed in the SIVP, and most existing sprouts underwent regression. Therefore, we defined the period following flow onset as the sprout regression phase. To determine whether this process depends on blood flow, we inhibited cardiac contraction and circulation using a morpholino targeting cardiac troponin T type 2a (*tnnt2a*) and examined SIVP morphology (Fig. 2). Although the overall size of the SIVP was unchanged in *tnnt2a* morphants compared to controls, (Fig. 2A-B’, C), its morphology was altered (Fig. 2A, B). Specifically, morphants exhibited fewer vascular loops (Fig. 2A’, B’ asterisks, D), reduced dorsoventral plexus length (Fig. 1A’, B’, E) and increased length (Fig. 2A’, B’, E) and number of leading sprouts (Fig. 2A”, B”, E, F). While the total EC number of the SIVP was not significantly different between morphants and controls (Fig. 2G), EC distribution was affected (Fig. 2A”, B”, H, I). Leading sprouts in morphants contained more ECs (Fig. 2A”, B”, H, I), whereas SIVP branches had significantly fewer. However, EC numbers in the ventral region of the SIVP (the sub intestinal vein, SIV), were not significantly altered (Fig. 2A”, B”, I).

**Figure 2.**
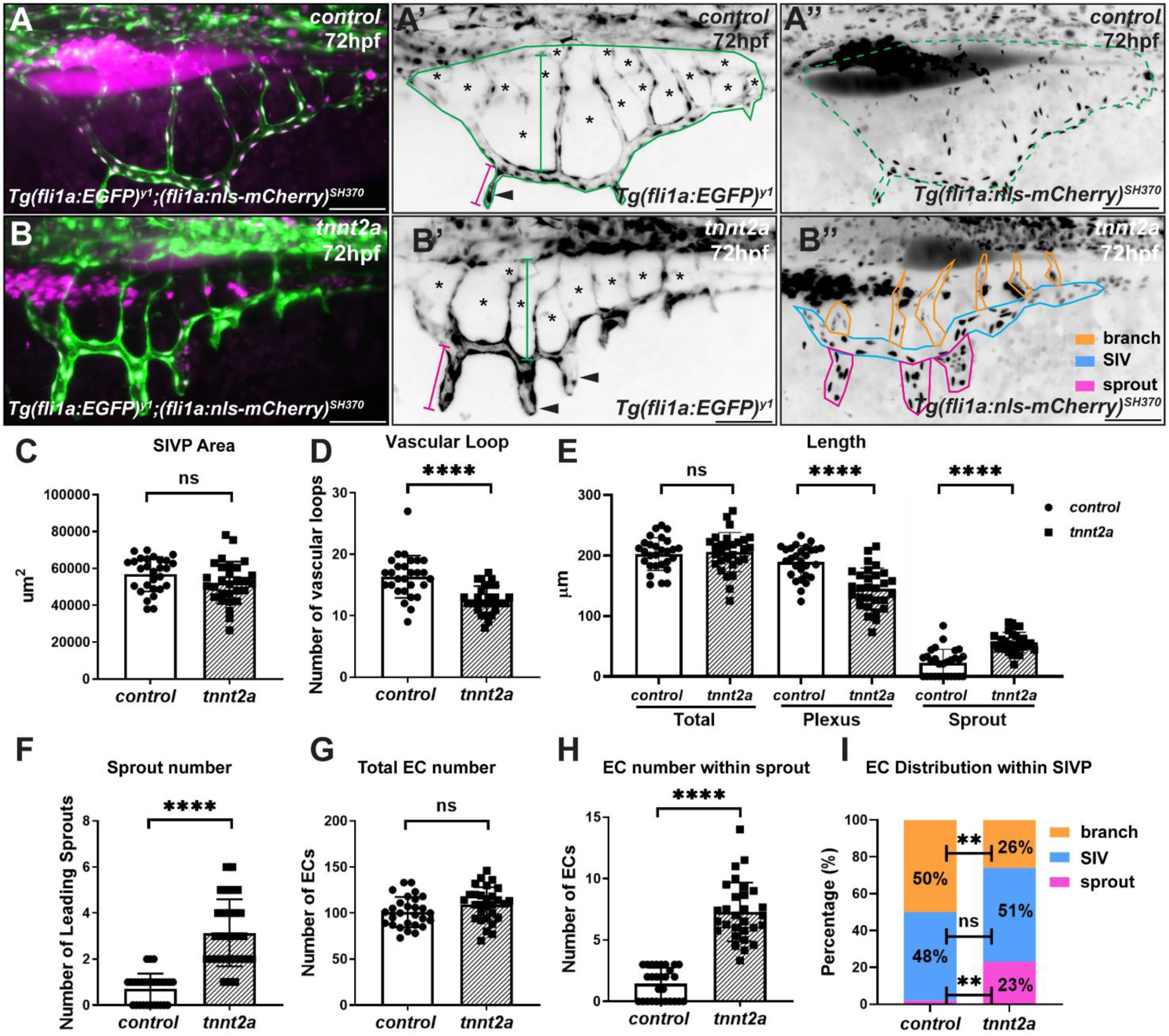
Blood flow controls morphology and endothelial cell distribution of the SIVP. **A**, **B**) Comparison of SIVP morphology at 72hpf in the presence (control morphants, **A**) or absence of blood flow (*tnnt2a* morphants). **B**) No significant difference in plexus area (highlighted in green in **A’**, **A’’**) was observed in the presence or absence of blood flow (**C**). Embryos without blood flow displayed fewer vascular loops (**A’**, asterisks, **D**). Total length of the SIVP plexus was not altered in the presence or absence of blood flow (**B**’, green line, **E**), however *tnnt2a* morphant embryos displayed reduced length of the SIVP basket (**B’**, blue line, **E**) and increased sprout length (**B’**, magenta line, **E**). The number of leading sprouts (**A’**, **B’** arrowheads) were increased in *tnnt2a* morphant embryos (**B”**, highlighted in magenta, **F**). The total number of ECs in the SIV was not altered by the flow status of the plexus (**G**), however, leading sprouts contained increased numbers of ECs in *tnnt2a* morphant embryos (**B”**, highlighted in magenta, **H**). In the absence of blood flow, SIV branches contained fewer ECs (**B”**, highlighted in orange, **I**). Unpaired *t*-test, \*\*\*\**p*≤0.0001, ns; *p*≥0.05, control embryos, n=28; *tnnt2a* embryos, n=29. Scale bars 50µm.

To control for potential off target effects induced by morpholino injection, we generated G0 *tnnt2a* CRISPR mutants (Wu et al., 2018) and assessed SIVP morphology. Mosaic *tnnt2a* mutants displayed abnormal plexus morphology (Fig. S1A, B arrows, C-I), phenocopying the defects observed in morphants (Fig. 2), confirming that these changes were specific to *tnnt2a* inhibition.

Together, these findings suggest that blood flow influences EC distribution within the developing venous plexus without affecting the total EC number. The increased size and frequency of leading sprouts in *tnnt2a* morphants further suggest that blood flow promotes sprout regression. Consistent with this, leading sprouts regressed in the presence of flow, with their ECs integrating into the SIV (Fig. 3A-C, arrows, Supplementary Movie 3). In contrast, sprout regression was significantly reduced in *tnnt2a* morphants which retained more leading sprouts by 72 hpf. Anastomosis of leading sprouts was rare and occurred independently of blood flow (Fig. 3A, asterisk, G, H, Supplementary movie 3). Overall, these data indicate that blood flow contributes to SIVP remodelling, in part by promoting regression of leading sprouts.

**Figure 3.**
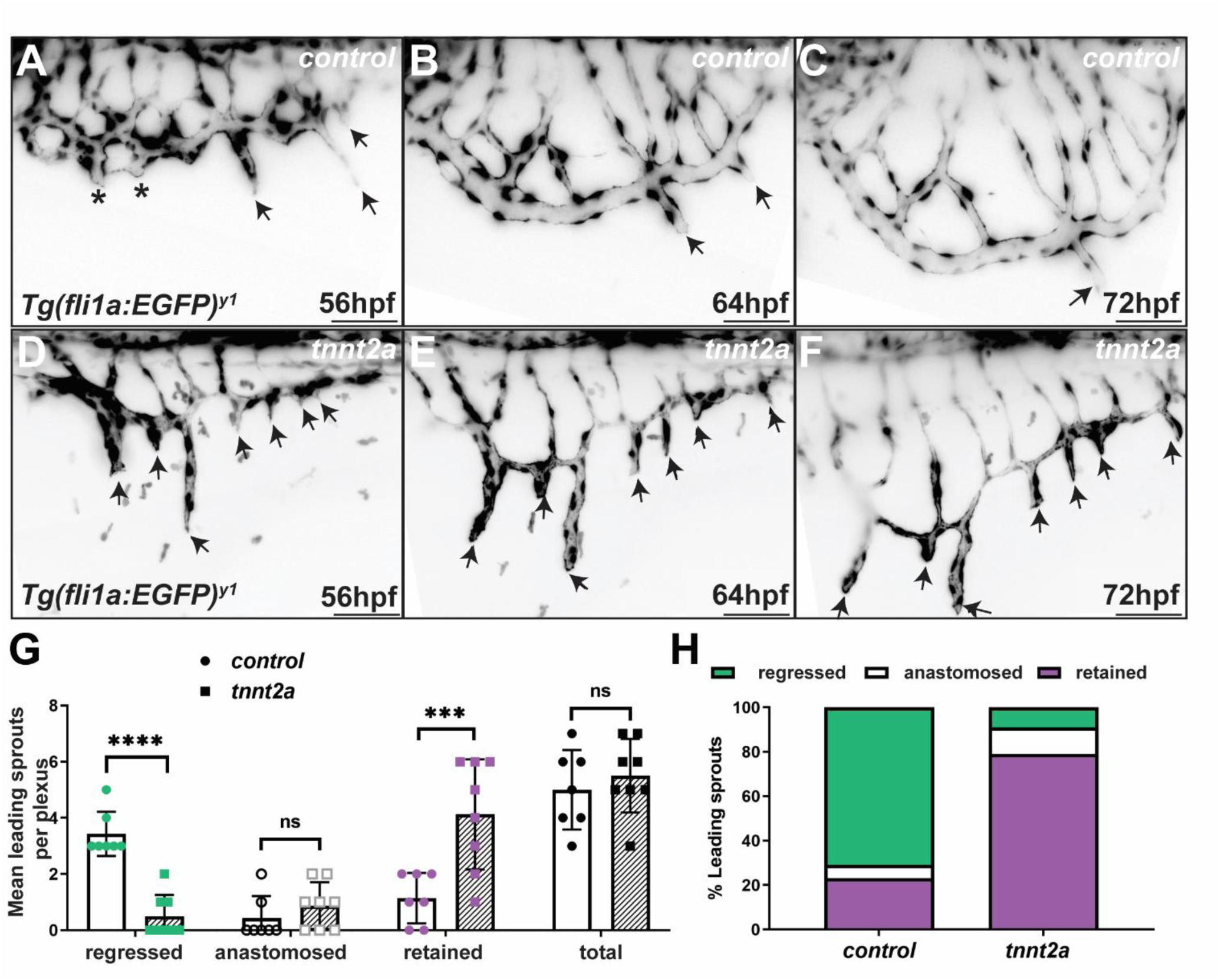
Blood flow promotes regression of SIVP leading sprouts. **A-F**) SIVP development 56-72hpf in the presence and absence of flow. Still images taken from time lapse movies (Supplementary Movie 3). **A**) In control animals, before blood flow enters the plexus, some angiogenic sprouts from the primary SIV anastomose (asterisks) to form vascular loops, while others lead at the migration front (arrows). **B**, **C**) In controls, leading sprouts regressed following the onset of blood flow and sprout ECs became incorporated into the SIV (arrows). **D-F**) In the absence of flow, leading sprouts failed to regress (**D-F**, arrows). **G, H**) Leading sprout regression was significantly reduced in *tnnt2a* morphant embryos and thus, the number of sprouts present at 72hpf were increased in comparison to controls. Unpaired *t*-test **** *p*≤0.0001; *** *p*≤0.001; ns *p*≥0.05; control morphants n=8; *tnnt2a* morphants n=7. Scale bars 50µm.

### Blood flow controls EC migration but not proliferation within the developing SIVP

The distribution of ECs within the developing SIVP could be influenced by several factors, including differences in cell proliferation, apoptosis, or migration. Although the total number of ECs within the SIVP remained unchanged in the absence of blood flow (Fig. 2G), the altered spatial distribution of ECs (Fig. 2H, I) suggested that flow might affect local rates of proliferation or apoptosis across. To investigate whether differences in EC proliferation, apoptosis, or migration contributed to the altered EC distribution observed in plexuses formed without blood flow (Fig. 2I), we tracked EC behaviour in the developing SIVP of control embryos and *tnnt2a* morphants between 56 and 70 hpf (Fig. 4, Supplementary Movie 4, 5). Similar numbers of ECs were tracked at the beginning of each time-lapse and ECs were grouped by their location at 56hpf, either within SIVP branches, SIV, or tip cells within leading sprouts (Fig. 4A-B). Endothelial proliferation was unaffected by the flow status of the developing plexus (Fig. 4C), and no apoptotic events were observed during SIVP development in either the presence or absence of blood flow. Thus, differential proliferation or apoptosis were unlikely to account for altered EC distribution observed in *tnnt2a* morphants (Fig. 2I). However, in the absence of blood flow, the frequency with which ECs differentially contributed to neighbouring EC subsets was altered (Fig. 4D). In the presence of blood flow, 73% of leading sprouts consisted of a pair of endothelial tip cells positioned in parallel to each other, which is consistent with previous reports (Hen et al., 2015). When tip cells divided in the presence of blood flow (Fig. 4B), the dorsal daughter cell rapidly migrated dorsally in most cases (77%), to contribute to the SIV (Supplementary Movie 6, arrows). This left the paired tip cells within the sprout, suggesting that the migrating daughter cell adopted a stalk identity (Supplementary Table 2). In rare cases (<2%) with blood flow present, daughter cells that migrated dorsally from leading sprouts to the SIV were later observed contributing to SIVP branches by 72 hpf (Fig. 4D). ECs were not seen migrating into leading sprouts under normal flow conditions. However, occasional bending and transient repositioning of EC nuclei at the boundary between the SIV and leading sprouts were observed, suggesting limited plasticity at this interface. In contrast, in the absence of blood flow, only 25% of tip cells migrated dorsally to the SIV, with most tip cells remaining within the leading sprout by 72 hpf. This aligns with the increased numbers of ECs observed in leading sprouts of *tnnt2a* morphants (Fig. 2B”, I, S1I). Notably, in *tnnt2a* morphants, SIV ECs were seen migrating ventrally into leading sprouts (Fig. 4D). Interestingly, these migrating ECs did not appear to compete with tip cells at the sprout front, suggesting they did not acquire tip cell identity.

**Figure 4.**
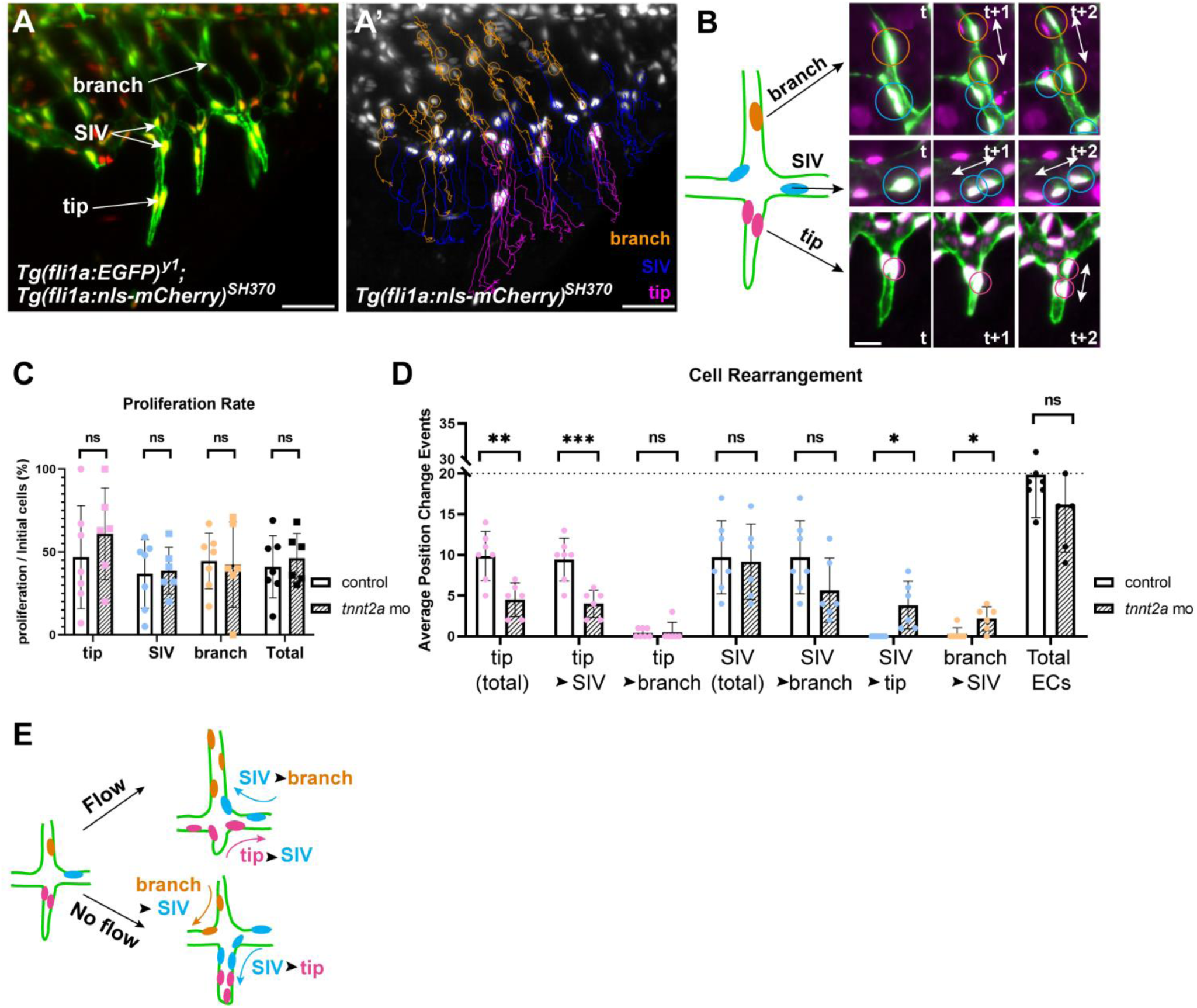
Blood flow coordinates EC migration but not proliferation within the developing SIVP. **A-B**) ECs in the SIVP were divided into three groups depending upon their initial position at 56hpf. SIVP branch cells (orange) within branches, SIV cells (magenta) within the ventral SIV (blue) and tip cells within leading sprouts (magenta). Examples of EC proliferation in subsequent timepoints are highlighted (**B**). The direction of cell division is parallel to the long axis of the vessel (**B**). **C**) EC proliferation rate (cell division events/initial number of ECs) did not differ in total or between groups in the presence or absence of flow. **D**) There was a substantial reduction in tip cell contribution to SIV in *tnnt2a* morphants, indicating that flow is required for dorsal tip cell rearrangement. In addition, there was an increase in ventral migration of SIV ECs into leading sprouts in the absence of flow. Most branch cells remained in the branches under flow, however, more branch ECs migrated ventrally to contribute to the SIV without blood flow. **E**) Schematic representation of differing EC migration in the presence and absence of blood flow. Unpaired *t*-test \*\*\**p*≤0.001; \*\**p*≤0.01; \**p*<0.05; ns *p*≥0.05, control morphants n=7; *tnnt2a* morphants n=6. Scale bars = 50µm (A-A’), 25µm (B).

ECs within SIVP branches make up approximately half of the total EC population within the plexus, yet they rarely change position in the presence of blood flow. Fewer than 1% of ECs in branches contributed to cellular rearrangement during SIVP development (Supplementary Table 2, Fig. 4D). In contrast, ECs from branches were significantly more likely to migrate into the SIV in *tnnt2a* morphants (Fig. 4D). Although the overall number of EC positional changes was not significantly different between control and *tnnt2a* morphants (Figure 4D), the direction of migration varied. ECs tended to migrate dorsally in the presence of flow, while ventral migration was more common in plexuses that developed without flow, resulting in an increased tendency for ECs to adopt and maintain tip cell behaviour (Fig. 4D). Collectively, these data suggest that blood flow is required to guide the directional rearrangement of ECs during SIVP development.

In the presence of blood flow, ECs within the SIVP exhibited distinct migratory behaviours depending upon their location. While the plexus expanded and ECs migrated ventrally, blood flow promoted dorsal migration, likely supporting coordinated plexus growth (Fig. 4E). In the absence of blood flow, ECs showed persistent ventral movement, leading to distended SIVP branches with fewer cells (Fig. 3C, F) and elongated leading sprouts with accumulated ECs displaying tip cell behaviour (Fig. 4E).

### Blood flow co-ordinates the direction of EC migration during SIVP development

During SIVP remodelling, ECs displayed distinct migratory behaviours depending upon their position within the plexus and the presence or absence of blood flow (Fig. 4D). To investigate whether blood flow influences EC migration, we focused on the most ventral ECs within each SIVP, particularly those located in the SIV and leading sprouts. Given the semi-circular curvature of the SIVP, ECs in the anterior and posterior of the SIVP migrate shorter distances than ventral ECs in the central region of the plexus. We tracked ECs from 56 hpf onward and analysed their movement between consecutive timepoints, referred to as migration steps.

ECs near the most ventral sprout were classified as either SIV ECs (blue) or tip cells (magenta) based on their initial position at 56 hpf (Fig. 5A, B), and migration steps were recorded at 30-minute intervals. While the total distance migrated by ECs did not differ significantly between controls and *tnnt2a* morphants (Fig. 5C), the meandering index (total distance/displacement ratio), was significantly reduced in tip cells in *tnnt2a* morphants (Fig. 5D). A meandering index closer to 1, indicates a straighter migration path, while values >1 reflect more tortuous paths. Thus, in the absence of blood flow, tip cells migrated more directly in a ventral direction, with reduced dorsal rearrangement toward the SIV (Fig. 5D). In contrast, the meandering index of ECs within the SIV was unaffected by flow status (Fig. 5E). Migration velocity of both tip and SIV ECs gradually decreased in the presence of blood flow, whereas no significant reduction was observed in *tnnt2a* morphants, (Fig. 5F), suggesting that flow may act to slow EC migration and promote vessel stability. Additionally, EC nuclear spacing was reduced in *tnnt2a* morphants (Fig. 5G), consistent with increased EC density in the absence of flow (Fig. 2B”).

**Figure 5.**
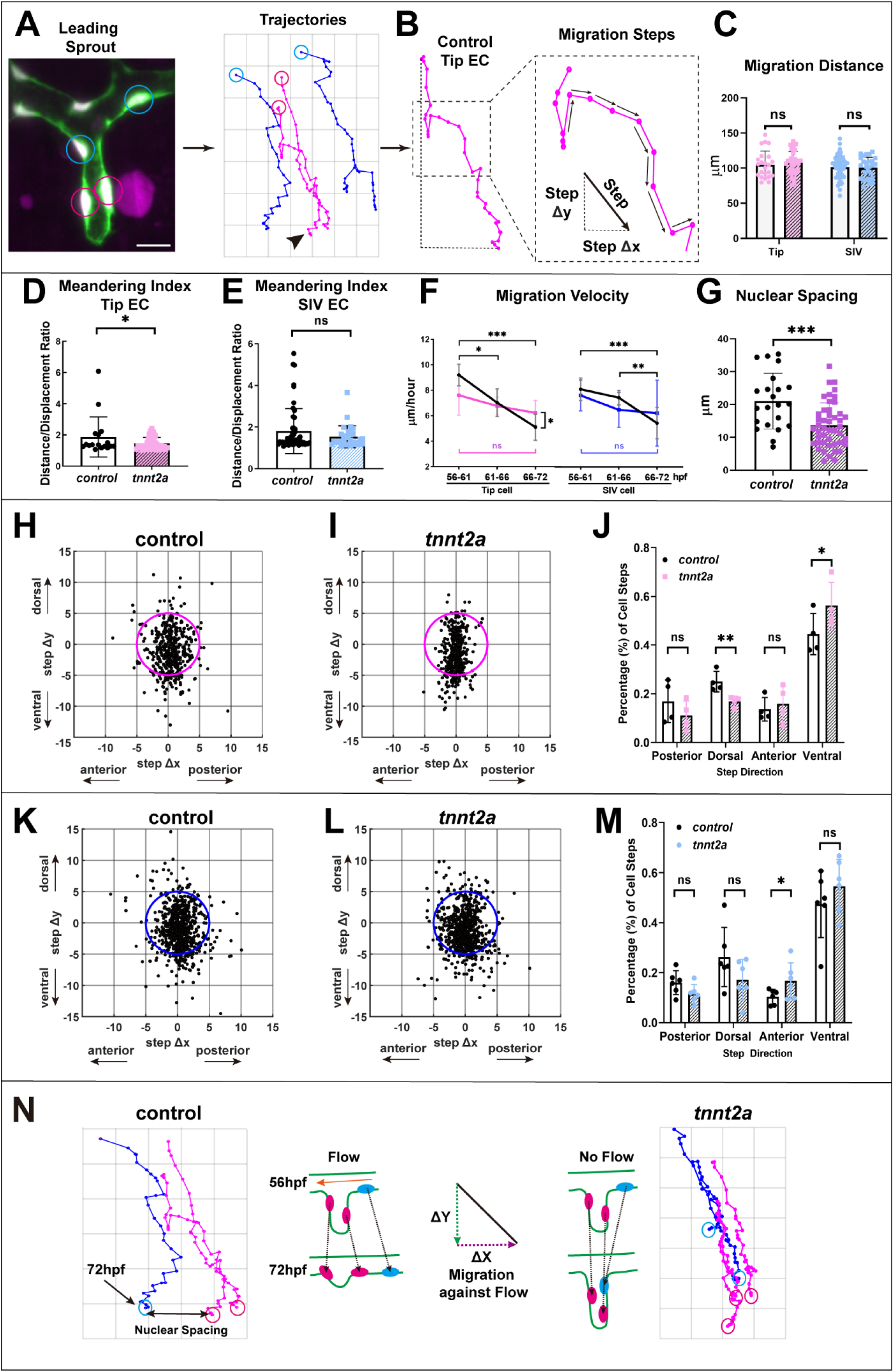
Blood flow controls EC migration direction during leading sprout regression. **A**) Examples of ECs within the SIV (blue) or tip cells (magenta) tracked in proximity to leading sprout and example migration trajectories. **B**) Migration ‘steps’ are defined as the movement of ECs between two consecutive time points (arrows). **C**) Migration distance (the sum of step displacements) of SIV ECs or tip cells was unaffected by flow status of SIVP. (**C-E**) Each dot represents a migration track of a single EC. control, n=79 tracks from 5 embryos; *tnnt2a*, n=64 tracks from 5 embryos. Tip cells: control, n=23 tracks, *tnnt2a*, n=37 tracks; SIV cells, control, n=56 tracks, *tnnt2a*, n=27 tracks; Unpaired *t*-test **D, E**) Meandering index was significantly reduced in tip cells in *tnnt2a* morphants (**D**) but was not significantly different in SIV ECs (**E**). **F**) Migration velocity of tip or SIV cells gradually reduced in the presence of flow (black lines) but was not significantly altered in *tnnt2a* morphants (coloured lines). Average speed of every step of tip cells or SIV cells from 6 control or *tnnt2a* embryos. Unpaired *t*-test. **G**) Nuclear spacing was significantly reduced in *tnnt2a* morphants indicating increased EC crowding within the SIVP. Control, n=22 from 6 embryos; *tnnt2a*, n=46 from 6 embryos. Unpaired *t*-test **H**-**J**) Scatter plots displaying coordinates of tip cell migration steps. The magenta circle in each plot (5µm radius) is outlined as a reference for coordinate distribution. Migration steps show increased alignment along Y-axis in *tnnt2a* mutants (**I**) in comparison to controls (**H**) and tip cells display significant reduction in dorsal and increase in ventral migration in the absence of blood flow (**J**). control, n=383 steps from 4 embryos; *tnnt2a*, n=860 steps from 4 embryos. **K-M**) Scatter plots displaying coordinates of SIV EC migration steps indicate increased migration steps in quadrant III in *tnnt2a* morphants (**L**) indicating anterior migration bias in the absence of blood flow (**M**). control, n=1313 steps from 6 embryos; *tnnt2a*, n=744 steps from 6 embryos. **N**) Schematic representation of EC migration within the SIVP during sprout regression. In the presence of flow, SIV cells migrate laterally to accommodate dorsally migrating tip cells as leading sprouts regress. In the absence of flow, SIV ECs migrate more anteriorly, and tip cells migrate ventrally leading to elongated sprouts and regression failure. **** *p*≤0.0001; *** *p*≤0.001; ** *p*≤0.01; * *p*<0.05; ns *p*≥0.05. Scale bar 25µm.

We defined migration steps as a relative change in position in both X and Y directions over a 30-minute interval (Fig. 5B). We plotted the overall change in position from their initial starting location at 56 hpf and found that most cells, whether SIV or tip cell, remained within a 5µm radius (Fig.5H, I, K, L). Tip cell migration displayed a larger difference between flow (control morphant) and no-flow (*tnnt2a* morphant) conditions, with control morphants displaying increased dorsal migration (Fig. 5H, J), and *tnnt2a* morphants displaying increased ventral migration (Fig. 5I, J), when compared to each other. Tip cell migration in morphants was more confined along the y-axis (Fig 5H, I, magenta circle), which reflects increased crowding within sprouts (Fig. 5G) and a shift toward ventral rather than dorsal migration (Fig. 5J). Migration steps for SIV cells displayed a markedly reduced difference between conditions (Fig. 5K, L). Most SIV EC migration steps fell within a 5µm radius, with control morphant SIV cells displaying a tendency towards posterior migration (Fig. 5K, M), while *tnnt2a* morphant SIV ECs displayed a significant anterior migration tendency (Fig. 5L, M). This suggests close contact between tip and SIV cells and collective migration of these during SIV development. However, in *tnnt2a* morphants, SIV ECs showed a significant anterior migration bias (Fig. 5K, L, M).

Together, these data suggest that blood flow provides directional cues for EC migration. During SIVP development, ECs collectively migrate in a ventral-anterior direction as the plexus expands. The onset of flow biases migration; SIV ECs migrate posteriorly, against the direction of flow, while tip cells migrate dorsally, leading to sprout regression and incorporation into the SIV (Fig. 5K). In the presence of flow, SIV ECs shift laterally to accommodate dorsally migrating tip cells as leading sprouts regress. In contrast, without flow, SIV ECs migrate anteriorly, and tip cells migrate ventrally, resulting in elongated sprouts and failed regression. Given that blood flows from posterior to anterior along the SIV (Fig. 1), these findings suggests that flow provides a horizontal mechanical force that guides EC migration during plexus remodelling.

### Blood flow polarises ECs to initiate reverse migration during sprout regression

During vascular development, ECs polarise in response to migration cues and blood flow (Franco et al., 2015; Kwon et al., 2016). Venous ECs typically exhibit lower levels of polarity compared to arterial ECs (Kwon et al., 2016). To determine whether ECs in the SIVP become polarised in response to flow during sprout regression, we used transgenic zebrafish lines *Tg(fli1a:nls-EGFP)^SH549^* and *Tg(fli1a:golgi-tagRFP, cryaa:CFP)^SH529^* to label endothelial nuclei and Golgi respectively (Fig. S2A-B). We imaged the SIVP in control and *tnnt2a* morphants at the onset of flow (56hpf) and during sprout regression (64 hpf) quantifying the relative positions of nuclei and Golgi in ECs of leading sprouts and the SIV.

Prior to flow onset, most tip cells showed ventral Golgi positioning at the migration front with a minority displaying dorsal polarity (Fig. S2B-C). This distribution was not significantly altered in *tnnt2a* morphants (Fig. S2C). Similarly, most SIV ECs were non-polarised before flow began, and no significant differences in polarity were observed between controls and morphants (Fig. S2D). By 64 hpf, once the SIVP was perfused, the proportion of tip cells with dorsally positioned Golgi was significantly reduced in *tnnt2a* morphants. This aligns with our observation that tip cells in morphants undergo persistent ventral migration whereas in controls, they tend to migrate dorsally (Fig. 5J). Additionally, by 64 hpf, a significantly higher proportion of SIV ECs in controls were polarised upstream of flow compared to morphants (Fig. S2B, D). These findings support our cell tracking data (Fig. 5) and suggest that blood flow provides directional cues that coordinate EC and initiate sprout regression.

To further explore EC rearrangement during sprout regression, we visualised the endothelial actin cytoskeleton using an EC-specific actin nanobody line, *Tg(fli1a:AC-TagRFP)* (Savage et al., 2019) (Fig. S3 & Supplementary Movie 7). Leading sprouts consisted of paired tip cells with their dorsal membranes connected to the SIV lumen (Fig. S3A-D). As regression progressed, both tip cells migrated over each other toward the SIV. The leading tip cell (Fig. S3E-H, green) reoriented its nucleus from a dorsoventral to lateral orientation, with most of its membrane incorporated into the perfused SIV. The adjacent tip cell (Fig. S3I-L, pink) followed, retracting its membrane and re-orienting its nucleus horizontally upon reaching the SIV. Meanwhile the leading tip cell (green) maintained its position, possibly to preserve vessel integrity. During this process, an F-actin positive focus formed at the ventral membrane of the trailing tip cell (Fig. S3H), which developed into a ring-like structure that expanded and eventually closed as the membrane merged with the main vessel (Fig. S3M-P, pink). This suggests dynamic actin reorganisation plays a role in sprout regression. Collectively, these observations indicate that exposure of the dorsal membranes of sprout ECs to blood flow within the SIV lumen is required for plexus remodelling via sprout regression initiation. Furthermore, the coordinated behaviour of neighbouring tip cells appears to ensure sprout regression without compromising vessel integrity.

### Leading sprout regression occurs under conditions of reduced blood flow

EC rearrangement from regions of low to high shear stress has been implicated in vessel pruning during vascular development (Franco et al., 2015). Shear stress experienced by ECs is directly influenced by haemodynamic flow and blood viscosity, and inversely proportional to the cube of the vessel radius (Heil and Schaper, 2004). Since leading sprouts in the SIVP are not perfused, and tip cells typically regress dorsally in response to SIV perfusion (Fig. 1H-J), we next asked whether reduced blood viscosity is compatible with normal sprout regression. To reduce blood viscosity, we titrated *gata1a* morpholino to deplete circulating erythrocytes by an average of 87% (Fig. 6A-C, arrowheads). Unlike *tnnt2a* loss-of-function (Fig. 2, S1), *gata1a* morphants showed no major changes in overall SIVP morphology (Fig. 6A, B). Interconnecting vessels (ICVs) remained connected to the SIA, and the number of vascular loops was unchanged (Fig. 6D). However, many ICVs were not patent in *gata1a* morphants (Fig. 6B), likely due to reduced erythrocyte flow which may have contributed to the smaller plexus size observed in these larvae (Fig. 6E, F). Surprisingly, the number of leading sprouts was not significantly different between control and *gata1a* morphants (Fig. 6G), indicating that sprout regression can occur under conditions of very low blood flow. In arteries, low shear stress generally induces inward remodelling to increase flow by constricting vessel diameter (Silver and Vita, 2006). These findings support the idea that SIVP remodelling via sprout regression can proceed under low-flow conditions and align with models proposing a low endothelial shear stress set point (Baeyens et al., 2015).

**Figure 6.**
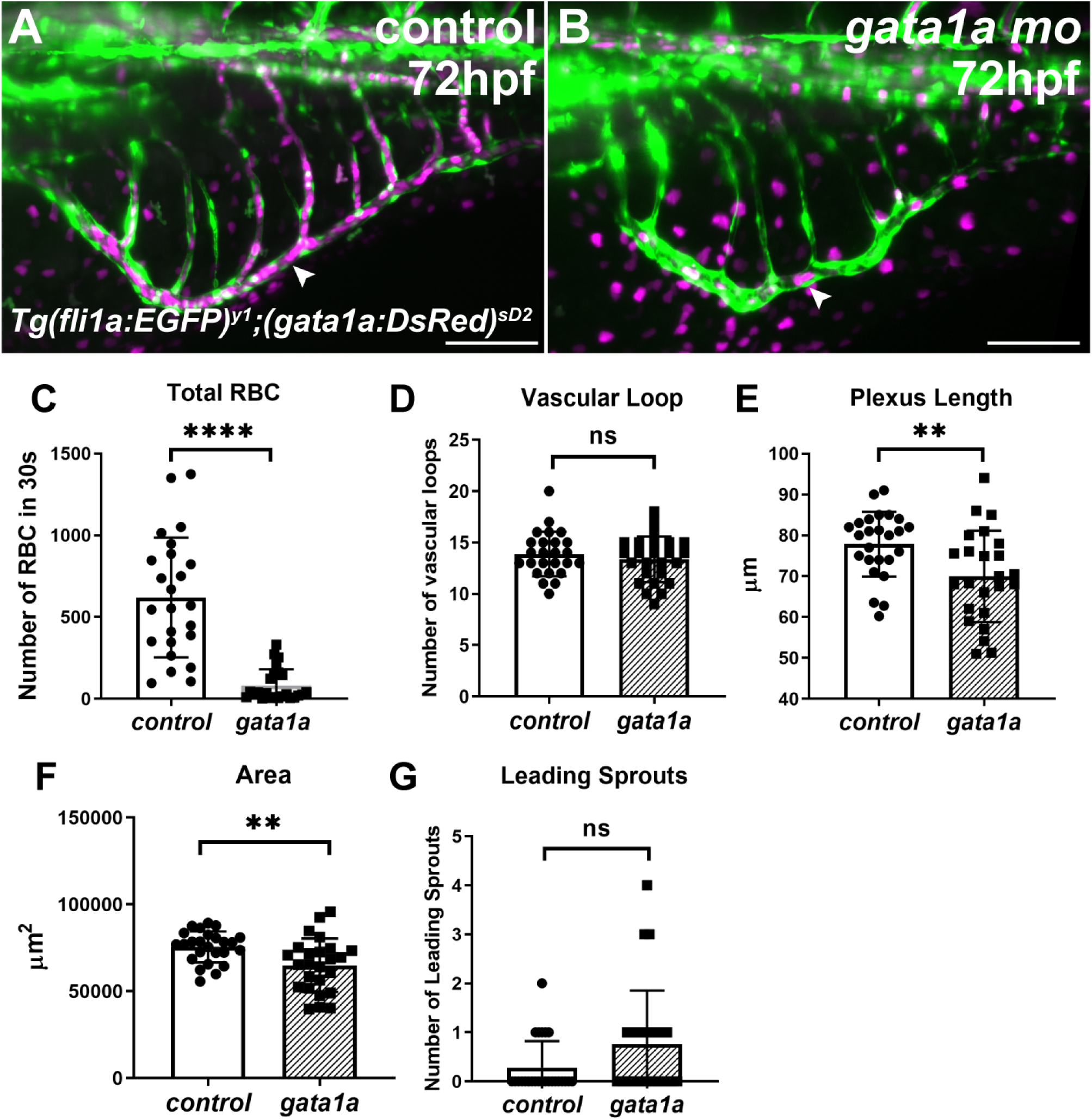
Leading sprout regression occurs under conditions of reduced blood flow. Leading sprouts undergo normal regression in *gata1a* morphants (**A**) in comparison to controls (**B**) despite a substantial reduction of circulating erythrocytes in *gata1a* morphants (**C**). Frequency of vascular loops did not differ between *gata1a* morphants and controls (**D**), but plexus length (**E**) and area (**F**) was reduced in *gata1a* morphants. Frequency of leading sprouts was not significantly altered in *gata1a* morphants (**G**). Unpaired *t*-test, **** *p*≤0.0001; ** *p*≤0.01; * *p*<0.05; ns *p*≥0.05; control morphants n=24, *gata1a* morphants n=24, 3 replicates. Scale bars 50µm.

### Leading sprout regression is dependent on *flt4* under low blood flow conditions

Zebrafish *vegfr4*/*kdrl* and *vegfr2*/*kdr* are individually and co-operatively required for normal SIV formation (Habeck et al., 2002; Koenig et al., 2016). In arteries, sustained deviations from the shear stress set-point trigger vascular remodelling via *flt4* in zebrafish arteries (Baeyens et al., 2015). Additionally, VEGFR1 (Flt1), inhibits excessive venous sprouting in the CNS (Krueger et al., 2011; Matsuoka et al., 2016; Wild et al., 2017) and may act as a decoy receptor to limit angiogenesis (Hiratsuka et al., 1998; Park et al., 1994; Zygmunt et al., 2011). Increased sprouting in the SIVP have previously been reported in *flt1* morphants (Hen et al., 2015), however, the flow status of the SIVP was not examined. We hypothesised that blood flow may regulate *flt1* expression in the SIVP to facilitate regression.

To test this, we first examined VEGF receptor expression in the SIVP under normal and flow-deficient conditions. Membrane-bound *flt1* (*mflt1*) was undetectable in the SIVP at 56 hpf when sprout regression typically begins (Fig. S4A, B), and at 72 hpf, when regression is complete (Fig S4C, D), regardless of flow status. Similarly, soluble *flt1* (*sflt1*) was not detected in the SIVP at 56 hpf in either condition (Fig. S4E, F). However, at 72 hpf, *sflt1* was present in the SIVP of 40% of *tnnt2a* morphants (Fig. S4G, H, arrow), but absent in controls, suggesting that blood flow may suppress *flt1* expression in the SIVP. This pattern was consistent in the cerebral vasculature where *sflt1* expression was present and unaffected by flow at 56hpf (Fig. S4I, J, arrows), but increased by 72hpf in morphants (Fig. S4K, L, arrows). Since *flt1* was not expressed in the SIVP under normal flow conditions, it is unlikely to be required for sprout regression. Supporting this, sprout regression was normal in *flt1* mutants (Fig. S5A-F) indicating *flt1* is dispensable for this process.

Kdrl is essential for proper formation and perfusion of the SIVP (Habeck et al., 2002), while Vegfr3/Flt4 has been proposed to act as a mechanosensor that regulates the endothelial shear stress set point (Baeyens et al., 2015). Both *kdrl* and *flt4* were broadly expressed throughout the developing SIVP (Fig. S6A-L, arrows) and their expression persisted in leading sprouts which failed to regress in *tnnt2a* morphants (Fig. S6I-L, arrows). *kdrl* expression was unaffected by blood flow status at all examined stages (Fig. S6A, B, E, F, I, J, arrows), whereas *flt4* expression was elevated in the ventral SIVP of *tnnt2a* morphants at 72 hpf compared to controls (Fig. S6K, L, green arrows). In controls, *flt4* expression was reduced during normal sprout regression but remained detectable in non-regressing sprouts in morphants (Fig. S6K, L arrows). Because loss of *kdrl* disrupts SIVP perfusion entirely (Habeck et al., 2002; Koenig et al., 2016) we focused on *flt4* Morpholino knockdown of *flt4* did not impair sprout regression and was compatible with normal remodelling, similar to what we observed with *gata1a* morphants (Fig. S7A-E, arrows). However, simultaneous knockdown of *flt4* and *gata1a* resulted in incomplete sprout regression by 72 hpf compared to controls (Fig. S7D, arrows, E).

Given known discrepancies between morpholino and mutant phenotypes (Kok et al., 2015), we generated a new *flt4* allele, *flt4^qmc316^*, using CRISPR mutagenesis This allele contains a frameshift mutation in exon 3 at amino acid 83, resulting in a premature truncation at amino acid 94. The frameshift is caused by a 144 bp insertion and an 8 bp deletion (Fig. S8A).*flt4^qmc316^*mutants showed no gross morphological defects by 5 dpf (Fig. S8B-C) but lacked a thoracic duct and exhibited impaired craniofacial and meningeal lymphatic formation (Fig. S8D-F), consistent with previously described *flt4* mutants (Hogan et al., 2009b; Shin et al., 2016). Importantly, *flt4^qmc316^* mutants do not display embryonic oedema up to 5 dpf, allowing us to study the genetic loss of *flt4* without disrupting blood flow in the SIVP. To assess whether *flt4* mutation affects sprout regression, we injected control and titrated *gata1a* morpholinos into embryos from a *flt4^qmc316^* heterozygous incross. Under normal flow conditions there was no significant difference in leading sprout number between wild-type, heterozygous and mutants (Fig. 7A-D). However, *gata1a* knockdown in *flt4^qmc316^* significantly increased the number of non-regressing sprouts compared to heterozygous and wild type siblings (Fig. 7E-H). This indicates that *flt4* function is required for effective sprout regression under low flow conditions. These data indicate that while *flt4* is dispensable for sprout regression under normal flow, it is essential for coordinating regression of leading sprouts when flow is reduced. Together, these findings demonstrate that blood flow plays a central role in coordinating endothelial cell migration and sprout regression during vascular plexus formation (Figure 8), establishing a foundation for further exploration of flow-responsive mechanisms in vascular development.

**Figure 7.**
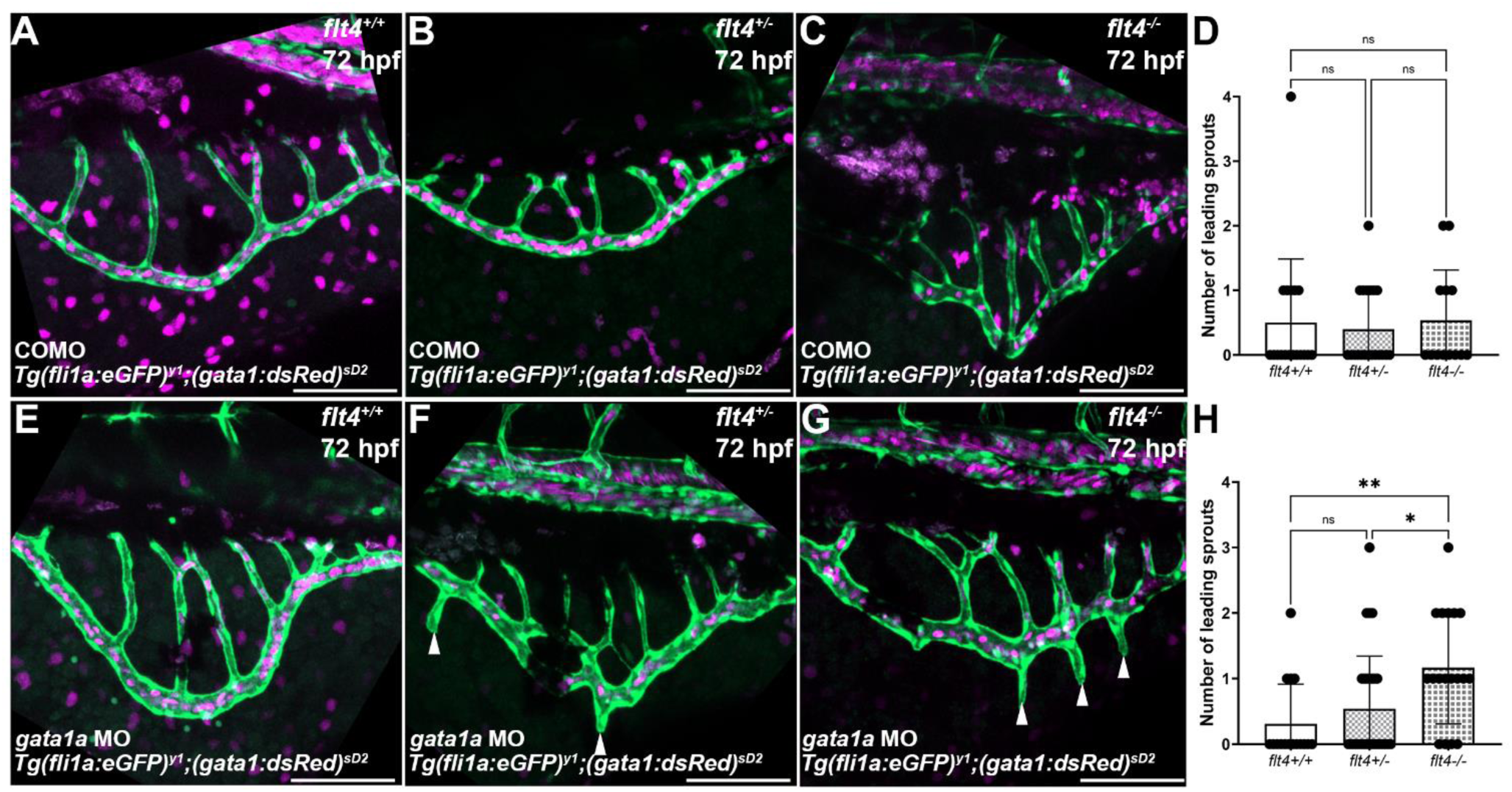
Leading sprout regression under low blood flow conditions is dependent on *flt4*. The number of leading sprouts observed in wild type (**A**), *flt4* heterozygous mutants (**B**), or *flt4* homozygous mutants (**C**) under normal flow conditions, injected with morpholino, was not significantly different (**D**). Low flow conditions, caused by *gata1* morpholino injection did not increase leading sprout numbers in wild type (**E)** or *flt4* heterozygous mutants (**F**), but leading sprouts in *flt4* homozygous mutants (**G**; white arrowheads) were significantly increased in number (**H**). One-way ANOVA, ** *p*≤ 0.01; * *p*≤0.05; ns *p*≥0.05. control morphant *flt4^+/+^* n=18; control morphant *flt4^+-+^* n=20; control morphant *flt4^-/-+^*n=13; gata1 morphant *flt4^+/+^* n=16; gata1 morphant *flt4^+-+^*n=37; gata1 morphant *flt4^-/-^* n=18, 3 replicates each. Scale bars 100µm.

**Figure 8.**
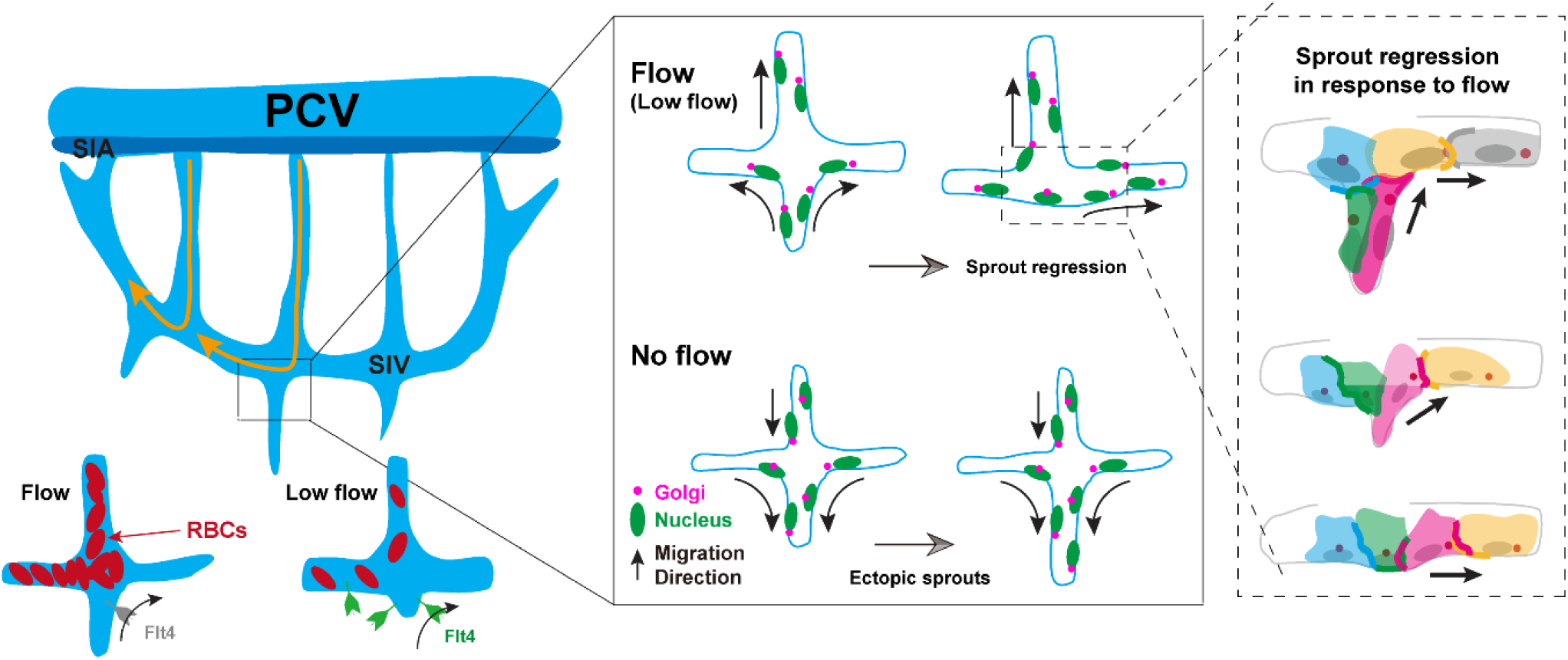
Cellular mechanisms of flow-mediated sprout regression. Schematic illustrations depicting the cellular behaviours during SIVP sprout regression in the presence and absence of flow. Blood flow promotes EC Golgi polarisation, dorsal and lateral EC migration and leads to sprout regression via Flt4 under low blood flow conditions. Illustrations highlighted with dashed lines depict co-ordination of EC migration during sprout regression in response to flow. ECs (in grey and yellow) at the luminal surface migrate laterally against flow and ECs (in green and pink) in the regressing sprout are pulled or use their neighbouring cells as migratory substrates via EC junctions (thickened lines), followed by EC rearrangements which lead to sprout regression. Abbreviations; PCV posterior cardinal vein; RBCs, red blood cells; SIV; sub-intestinal vein; SIA, supra-intestinal artery

## Discussion

Developmental remodelling of vascular networks has been described in different models and has mainly focused on pruning of vascular cross branches (Chen et al., 2012; Franco et al., 2015; Kochhan et al., 2013; Lobov et al., 2011; Phng et al., 2009). In this context, pruning occurs in loop forming or cross branch vessel segments, which ultimately leads to regression of the cross branch, leaving behind the two parallel vessels via a process involving EC migration and apoptosis (Chen et al., 2012; Zhang et al., 2018). Dynamic EC migration and rearrangement have also been described in sprouts, however studies have mostly focused on angiogenic sprouting and anastomosis (Arima et al., 2011; Bentley et al., 2014; Jakobsson et al., 2010). Whether sprouts ultimately anastomose with each other, or not, and how these sprouts are remodelled, remained unknown. Here using the zebrafish SIVP we find most leading sprouts within the developing plexus are remodelled via EC migration-driven regression rather than anastomosis. Apoptosis during pruning has been described in murine retinal vessels when circulation is compromised (Franco et al., 2016, 2015; Hughes and Chan-Ling, 2000) and also in zebrafish cranial arteries (Kochhan et al., 2013; Zhang et al., 2018) but this has not been implicated during branch pruning within the SIVP (Lenard et al., 2015). Consistent with this, we tracked approximately 70% of SIVP ECs per plexus and, while we cannot exclude the possibility that EC apoptosis occurs in the most anterior or posterior SIVP, or in the SIA, we observed no apoptotic events following live imaging of over 100 leading sprouts. Therefore, our data suggest that EC migration represents the primary mechanism of sprout regression within the SIVP. Indeed, angiogenic regression is an efficient system because cells are recycled via migration from sprouts, which are temporary structures, to other more permanent structures within developing vessels. Thus, co-ordination of EC migration represents a more efficient use of resources than sculpting vascular networks by proliferation or apoptosis.

Haemodynamic force exerted by blood flow has been proposed to be crucial in selecting and triggering pruning and regression of particular vessels (Chen et al., 2012; Franco et al., 2015; Kochhan et al., 2013; Lenard et al., 2015; Lucitti et al., 2007). Pruning mostly occurs in small and bifurcated branches with relatively unstable, or low, blood flow in comparison to adjacent large vessels (Chen et al., 2012; Lenard et al., 2015). This observation suggests that pruning is triggered by local differences in flow patterns between branches, which in turn induce EC polarisation and direct EC migration against flow, from low-flow to high flow vessels (Franco et al., 2015). Our data are consistent with this and upon initiation of sprout regression within the SIVP, ECs within the non-perfused leading sprouts are likely ‘attracted’ by the relatively high flow in the SIV, these polarise and migrate out of the regressing sprouts and subsequently contribute to the SIV. In line with this, dorsal-lateral polarisation and migration-induced EC rearrangement are highly dynamic processes during sprout regression and are dependent on blood flow. Interestingly, ventral EC migration persisted throughout the SIVP in *tnnt2a* morphants which indicates a flow-independent mechanism controls this aspect of EC migration. While blood flow co-ordinates remodelling of the SIVP during development, the plexus forms a stereotypical basket-like structure consisting of well-defined branches, SIV and leading sprouts even in the absence of flow. This suggests SIVP formation is regulated by well-orchestrated molecular patterning cues which may interact with physical forces such as blood flow to refine plexus development. Indeed, loss of guidance molecules such as PlexinD1 disrupt SIVP patterning (Goi and Childs, 2016) and PlexinD1 has recently been identified as a shear stress mechanosensor in ECs (Mehta et al., 2020), however, it is unclear whether plexinD1 modulates EC response to flow during SIVP remodelling.

Remodelling of blood vessels from a primitive structure into a more mature network at onset of flow allows temporal separation of sprouting at the migration front from pruning of branches during angiogenesis (Lenard et al., 2015). In the SIVP, new sprouts are rarely produced after the onset of flow and most existing sprouts remodel via regression. While the entire plexus expands ventrally during this period, the tip cells within leading sprouts do not simply undergo initial ventral migration followed by dorsal migration during regression, rather these ECs display transient and stochastic dorsal migration steps while undergoing collective, flow-independent, ventral migration. This suggests that sprouting and regression are interlinked. An intriguing question is at which point such steps lead to irreversible sprout regression. One possibility is when the tip cells in the sprout come into direct contact with blood flow. At the onset of flow, the posterior membranes of the tip cells are connected with adjacent SIV ECs and have limited exposure to blood flow within the SIV. It is possible that when blood flow creates sufficiently high shear forces on the SIV cells, it drives their migration against flow, and this may exert a pulling force on the tip cells which could promote their dorsal-lateral migration and induce sprout regression, via changes in polarisation. The SIV cells and the tip cells in contact with flow could act as the bridging ECs to sense differences in the local flow environment, as occurs during branch pruning (Franco et al., 2015). That the combined plexus and sprout length in *tnnt2a* morphants was not significantly different to controls indicates that in the absence of blood flow, ECs within sprouts can migrate the same distance as in controls. Reduced plexus length in the absence of blood flow is therefore a consequence of increased contribution of ECs from the branches and SIV to leading sprouts. In the presence of blood flow, ECs within sprouts migrate against flow and contribute to the SIV and branches, thereby supporting their expansion or growth. This is consistent with reports that ECs incorporate in developing arterial networks in the retina and coronary vasculature via coordinated EC migration (Chang et al., 2017; Pitulescu et al., 2017). Our data therefore suggests blood flow promotes regression of leading sprouts within the SIVP as a mechanism to support expansion of the developing plexus. Moreover, sprouting and regression share similar cellular behaviours but in a ‘reverse mode’.

Yolk sac vascular remodelling is dependent upon erythroblast circulation (Lucitti et al., 2007) and uneven erythrocyte distribution in bifurcations has been posited to enhance local shear stress differences between vessels undergoing pruning and their neighbouring vessels (Zhou et al., 2020). Thus, blood viscosity is strongly associated with vessel regression. How ECs sense low levels of flow remains unclear, but several possible molecular mechanisms have been proposed. The level of VEGFR3 has been suggested to maintain a threshold of shear stress sensed by ECs (Baeyens et al., 2015). Interaction of the transforming growth factor β (TGFβ) receptor, activin receptor like kinase ACVRL1/ALK1 with a component of the TGFβ receptor complex, Endoglin, has been reported to enhance EC sensitivity of flow sensing (Baeyens et al., 2016). In addition, non-canonical Wnt signalling may control vessel regression by modulating the threshold for flow-induced EC polarisation (Franco et al., 2016). We find that while plexus length was reduced in erythrocyte depleted embryos, even substantial reductions in blood viscosity were compatible with sprout regression. We find that sprout regression in the SIVP is dependent upon *flt4* under these reduced flow conditions, but *flt4* function is dispensable for sprout regression under normal blood flow. This indicates additional mechanisms could be involved to regulate sprout regression under normal flow conditions. Consistent with this, *flt4* mutants display normal SIVP formation (Hogan et al., 2009a; Hogan et al., 2009b), suggesting different functions of Flt4 under different flow conditions in zebrafish. The established role of VEGFR2 during EC mechanosensation (Coon et al., 2015; Tzima et al., 2005) suggests Kdr and Kdrl are promising candidates to mediate sprout regression under normal flow, however, their requirement during SIVP formation precluded their assessment in this study. The redundant function of Flt4 during sprout regression imparts robustness on the system and facilitates proper vessel remodelling under dysregulated flow conditions. It is important to note that while SIVP leading sprouts are not perfused before or during their regression, a low level of shear provided by skimmed plasma could promote dorsal migration of tip cells to elicit regression (Zhou et al., 2020). Future work will be needed to determine how *flt4* integrates with other mechanosensory pathways and how the mechanosensory input from altered blood flow is transduced by Flt4 to elicit the cytoskeletal changes necessary to rapidly adjust EC migration.

Together, our data demonstrates that blood flow drives remodelling of a venous plexus by coordinating EC migration and sprout regression and identifies *flt4* as a key mediator under low flow conditions. These insights highlight the dynamic interplay between mechanical forces and molecular signalling in shaping vascular architecture and underscore the importance of flow-responsive pathways in venous remodelling.

## Acknowledgements

We would like to thank the aquarium staff at the University of Sheffield and University of Nottingham for excellent zebrafish husbandry. We thank Freek van Eeden for helpful discussions, Didier Stainier for providing the *flt1* mutant and Katie Fisher for bioinformatic support with cell tracking. This work was supported by a China Scholarship Council and University of Sheffield joint studentship to YC, JG Graves Medical Research Fellowship and Nottingham Research Fellowship to RNW, Diabetes UK (17/0005678 to RNW), BBSRC (BB/R015457/1 to RNW), MRC (MR/X008215/1 to RNW) and British Heart Foundation (RG/19/10/34506 to PCE).

## Author Contribution

RNW and PCE conceived the project. YC, ZJ, HRK, AMS, PCE, RNW designed experiments; YC, ZJ, HRK, AMS performed experiments, YC, ZJ, AMS, RNW analysed data, RNW and AMS wrote the manuscript and all authors reviewed the final manuscript.

