## Supplementary Figures for "Blood flow coordinates collective endothelial cell migration during vascular plexus formation and promotes angiogenic sprout regression via *vegfr3/flt4*"

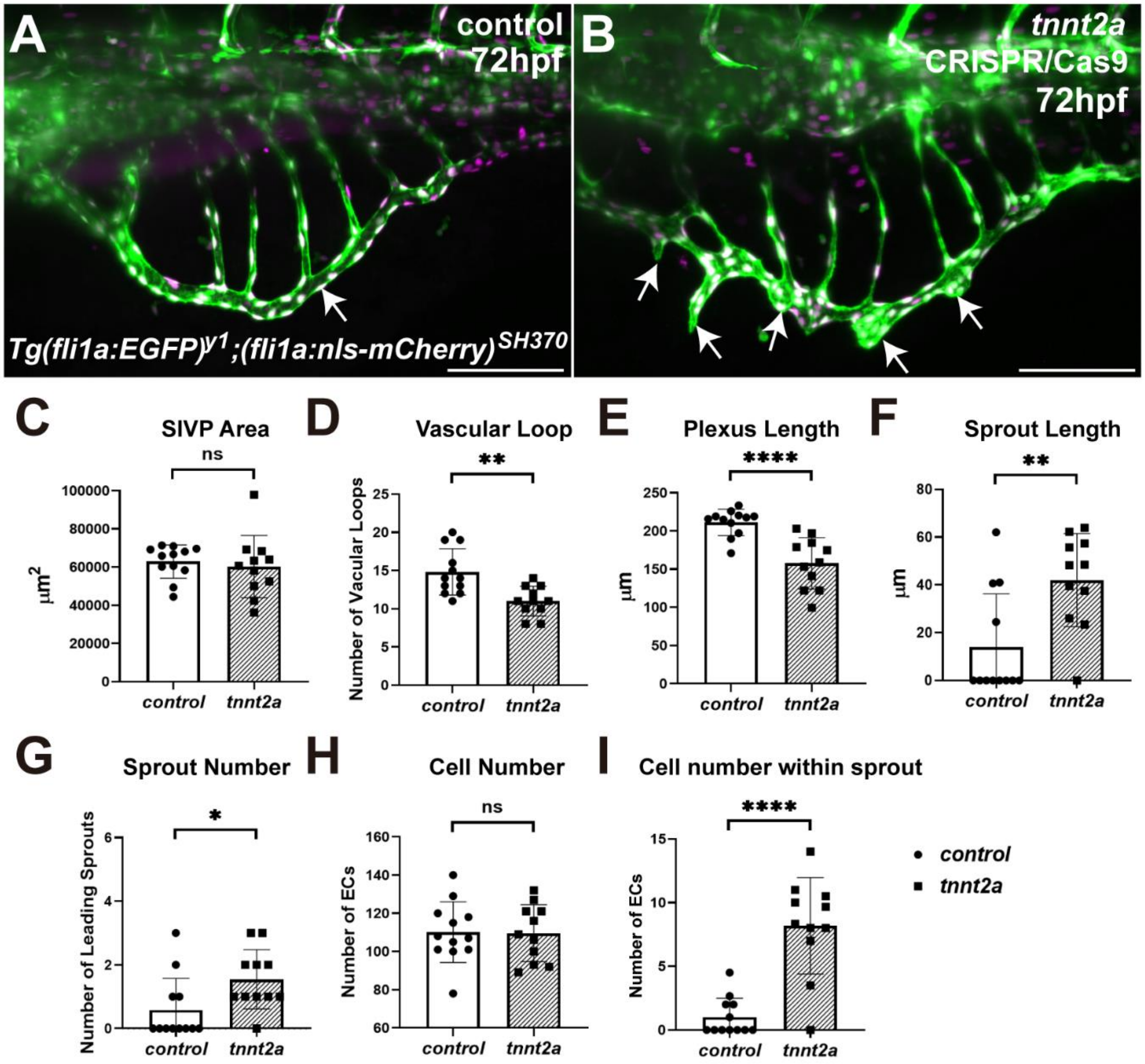


**Fig. S1 SIV development in *tnnt2a* F0 mosaic CRISPR mutants phenocopies *tnnt2a* morphants**

**A**, **B**) Embryos were injected with control gRNA and tracr RNA or *tnnt2a* gRNAs and tracr gRNA alongside nCas9 protein. Comparison of SIVP morphology at 72hpf in the presence (control, **A**) or absence of blood flow (*tnnt2a* mosaic mutant, **B**). **C**) No significant difference in plexus area was observed between control or *tnnt2a* mosaic mutants. **D, E**) *tnnt2a* mosaic mutants displayed fewer vascular loops than controls (**D**) and length of SIVP basket was reduced (**E**). **F**, **G**) *tnnt2a* mosaic mutant embryos displayed increased sprout length and number in comparison to controls. **H, I**) The total number of ECs in the SIV was not altered in *tnnt2a* mosaic mutants in comparison to controls, however, leading sprouts contained more ECs in *tnnt2a* mosaic mutants (**I**). Control, n=12; *tnnt2a*, n=11; Unpaired t-test, ****$p$≤=0.0001, ns, $p$≥0.05. Scale bars 100μm.

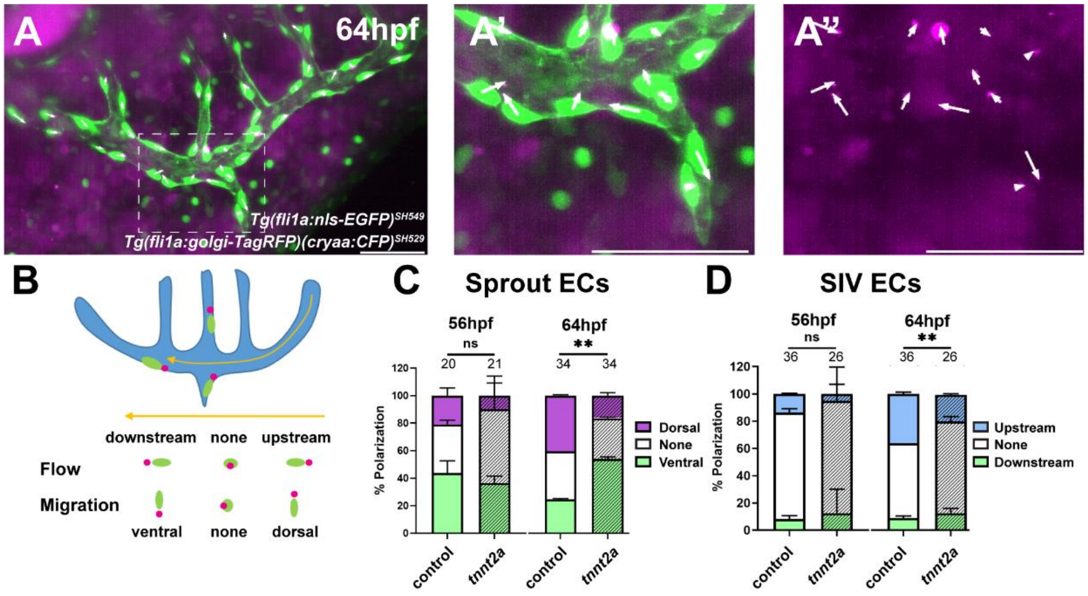


**Fig. S2 Blood flow promotes Golgi polarisation of ECs in the SIVP during sprout regression**
**A**) Examples of endothelial polarity based on relative position of nucleus and Golgi. Area highlighted in **A** is enlarged in **A'** *Tg(fli1a:nls-EGFP)* and **A''** *Tg(fli1a:golgi-TagRFP)*. Arrows illustrate Golgi orientation relative to nucleus. **B**) Schematic representation of EC polarity as indicated by relative nucleus-Golgi position. Arrows indicate the direction of blood flow. ECs polarise in the direction of, or in opposition to, blood flow. Golgi polarity was quantified at onset of flow (56hpf) and during sprout regression (64hpf) in ECs within leading sprouts or the SIV (**C** & **D**). **C, D**) Comparison of proportion of polarised ECs in leading sprouts in control and *tnnt2a* morphants at different time points. There was a significant decrease in the percentage of dorsally polarised sprout cells in the absence of flow at 64hpf (**C**, magenta) and in the percentage of upstream polarised SIV cells in the absence of flow at 64hpf (**D**, blue). Cell numbers are indicated above the bars. Unpaired t-test, *$p<0.05$, **$p\leq0.01$, ns, $p\geq0.05$. Scale bars 50µm.

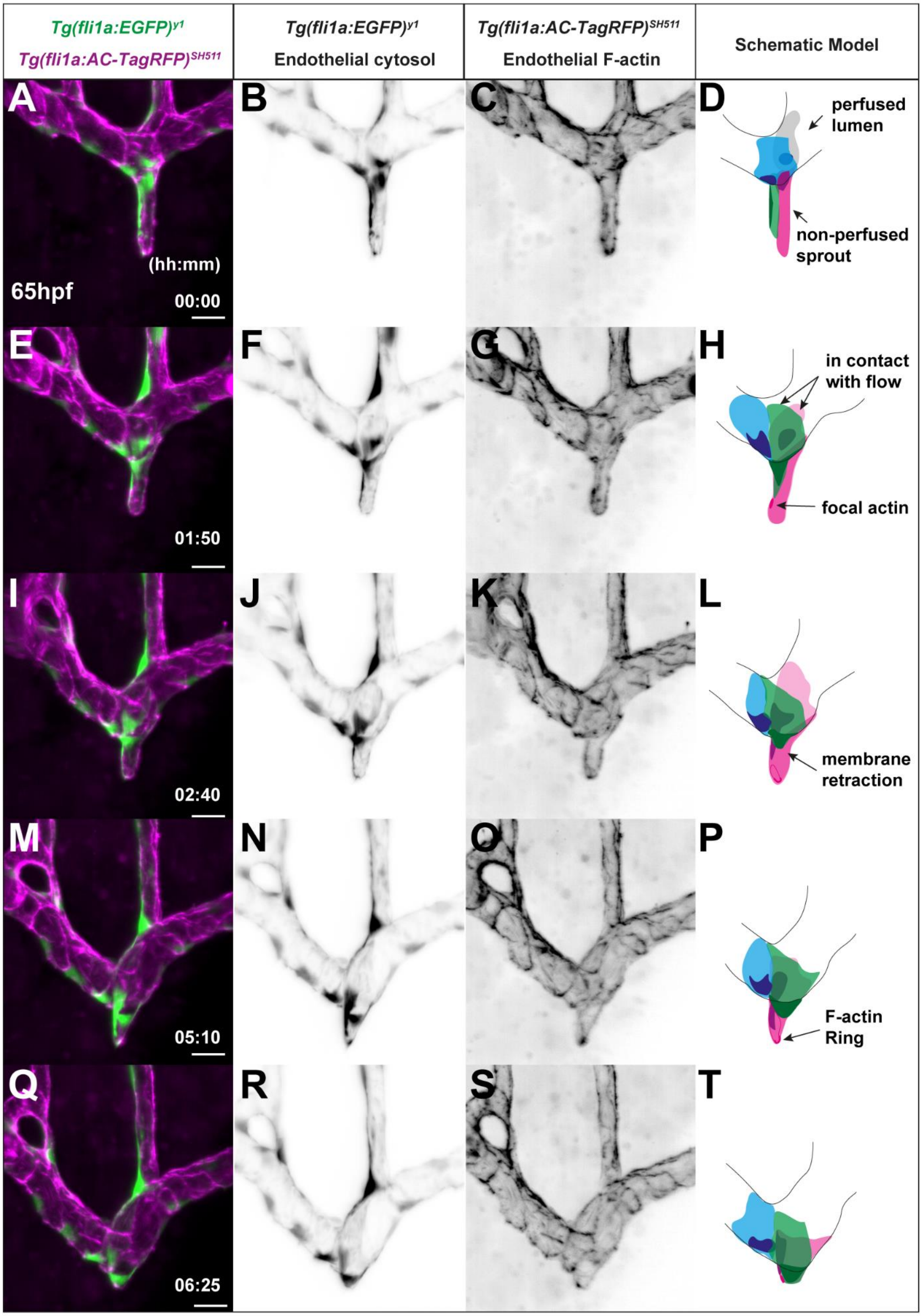

Tg(fli1a:EGFP)y1
Tg(fli1a:AC-TagRFP)SH511
Tg(fli1a:EGFP)y1
Endothelial cytosol
Tg(fli1a:AC-TagRFP)SH511
Endothelial F-actin
Schematic Model
A
B
C
D
perfused lumen
non-perfused sprout
(hh:mm)
65hpf
00:00
E
F
G
H
in contact with flow
focal actin
01:50
I
J
K
L
membrane retraction
02:40
M
N
O
P
F-actin Ring
05:10
Q
R
S
T
06:25

**Fig. S3 Cellular mechanisms of sprout regression**

(**A-E**) Representative images of leading sprout regression taken from 6.5h time-lapse from 65hpf in *Tg(fli1a:EGFP)*$^{y1}$*;(fli1a:AC-TagRFP)*$^{SH511}$ embryo with normal blood flow (Supplementary Movie 7). GFP (EC cytosol and nucleus) and RFP (EC actin) channels are shown in addition to merged image. Schematic model is drawn to illustrate the dynamic process of EC arrangement. **A-D**) Leading sprout consisting of a pair EC tip cells. The dorsal most EC membrane is in contact with blood flow within SIV lumen. **E-H**) During regression, one of the pair of tip cells (green) retracts most of its cell body within the SIV and positions its nucleus at the edge of the main vessel. The neighbouring tip cell (pink) retracts its membrane into the SIV while the distal region of the tip cell remains within the sprout. **I-L**) One of the tip cells (pink) continues to retract its distal membrane while the adjacent tip cell (green) holds its position and maintains vessel integrity. **M-P**) F-actin positive focus in the distal tip cell (pink) membrane grows into a ring which closes when the tip cell completes sprout regression. **Q-T**) Both tip cells complete regression and become part of the SIV. Scale bars 25μm.

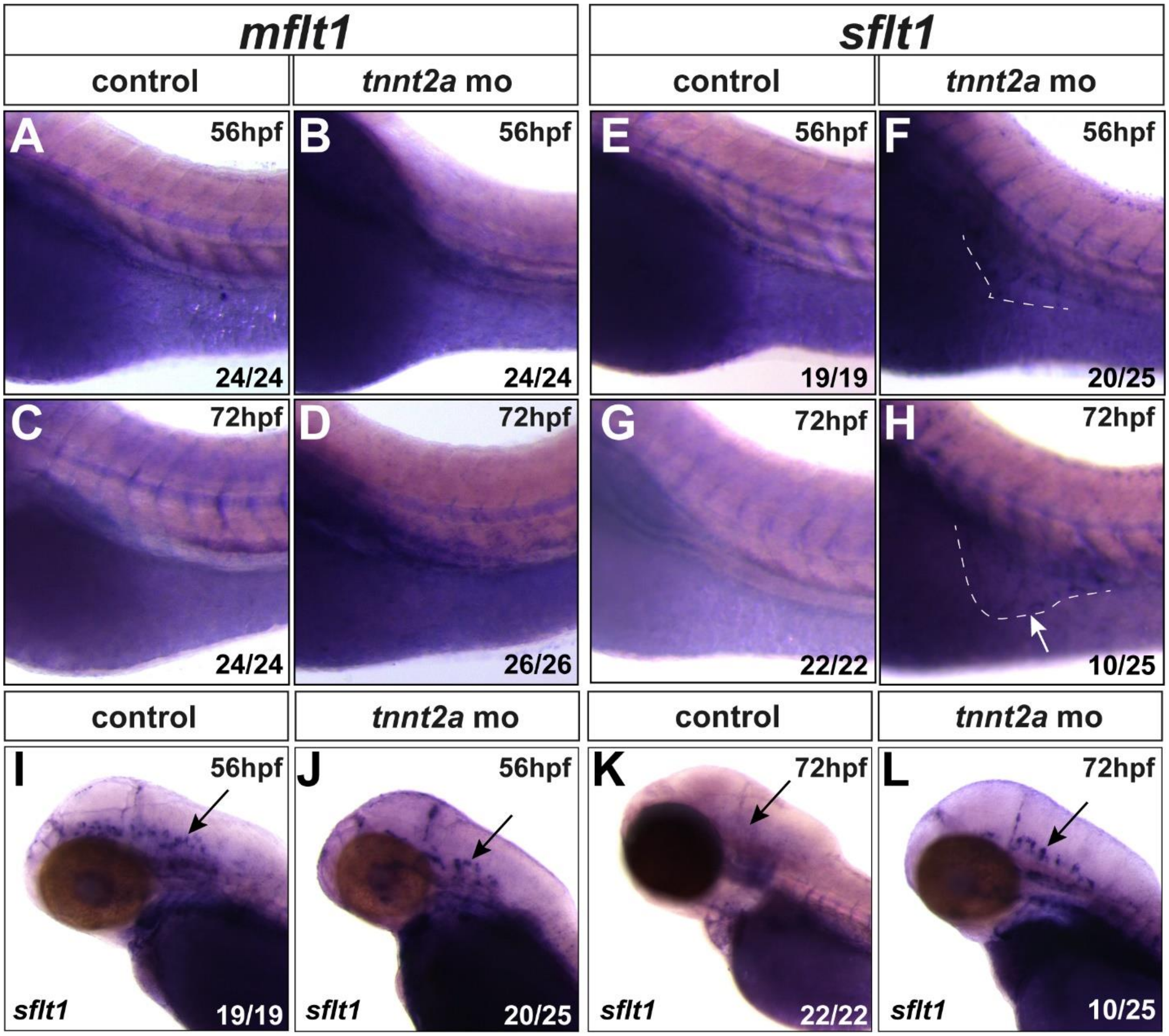


**Fig. S4 *sflt1* expression is increased in the SIVP in the absence of blood flow**

**A**-**D**) Expression of *mfl1* was not detected in the presence or absence of blood flow within the SIVP during the period of leading sprout regression. **E-H**) *sflt1* was not expressed in the SIVP in the presence of blood flow at the onset of leading sprout regression, but by 72hpf at the completion of regression, *slft1* was present in the SIVP only in the absence of blood flow (**H**, arrow). **I**, **J**) *sflt1* was expressed within central arteries at 56hpf in both the presence (**I**, arrow) and absence (**J**, arrow) of blood flow. **K**, **L**) slft1 expression was absent from central arteries by 72hpf (**K**, arrow), but persisted in embryos without flow (**L**, arrow).

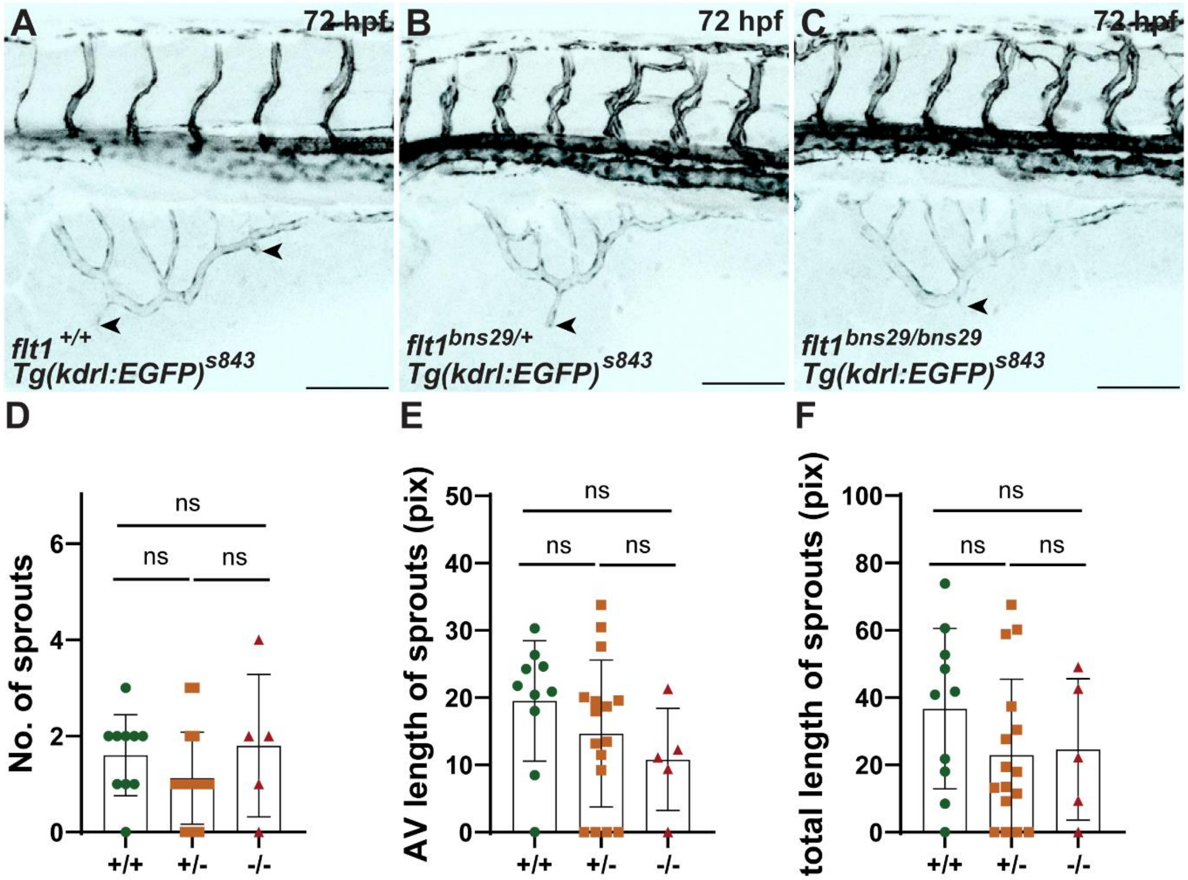


**Fig. S5 Leading sprout regression occurs normally in the SIVP of *flt1* mutants**

**A-F)** No significant differences were observed in the frequency (**D**), mean (**E**) or total length of sprouts measured in pixels (pix) (**F**) in WT, *flt1* heterozygotes or *flt1* mutant embryos at 72hpf. Unpaired t-test ns, $p \geq 0.05$. Scale bars 100µm.

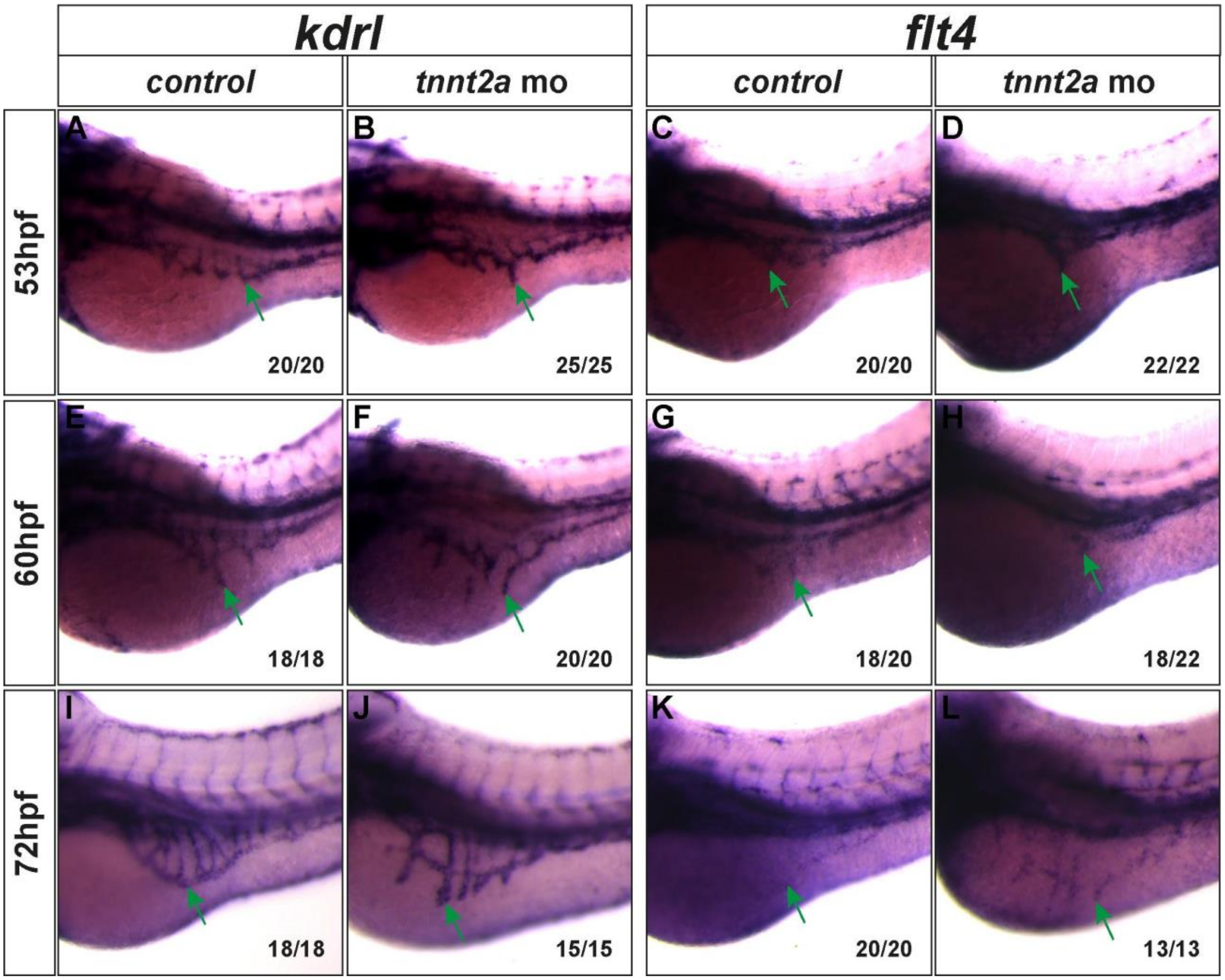


**Fig. S6 Expression of *vegfr4/kdrl* and *vegfr3/flt4* are retained in leading sprouts in *tnnt2a* morphants**

Both *vegfr4*/*kdrl* and vegfr3/*flt4* were expressed in ECs of the SIVP at before onset of flow at 53hpf, during sprout regression at 60hpf, and 72hpf in the presence or absence of flow. (green arrows). The expression of *kdrl* and *flt4* were retained in leading sprouts which had failed to regress in the absence of flow by 72hpf.

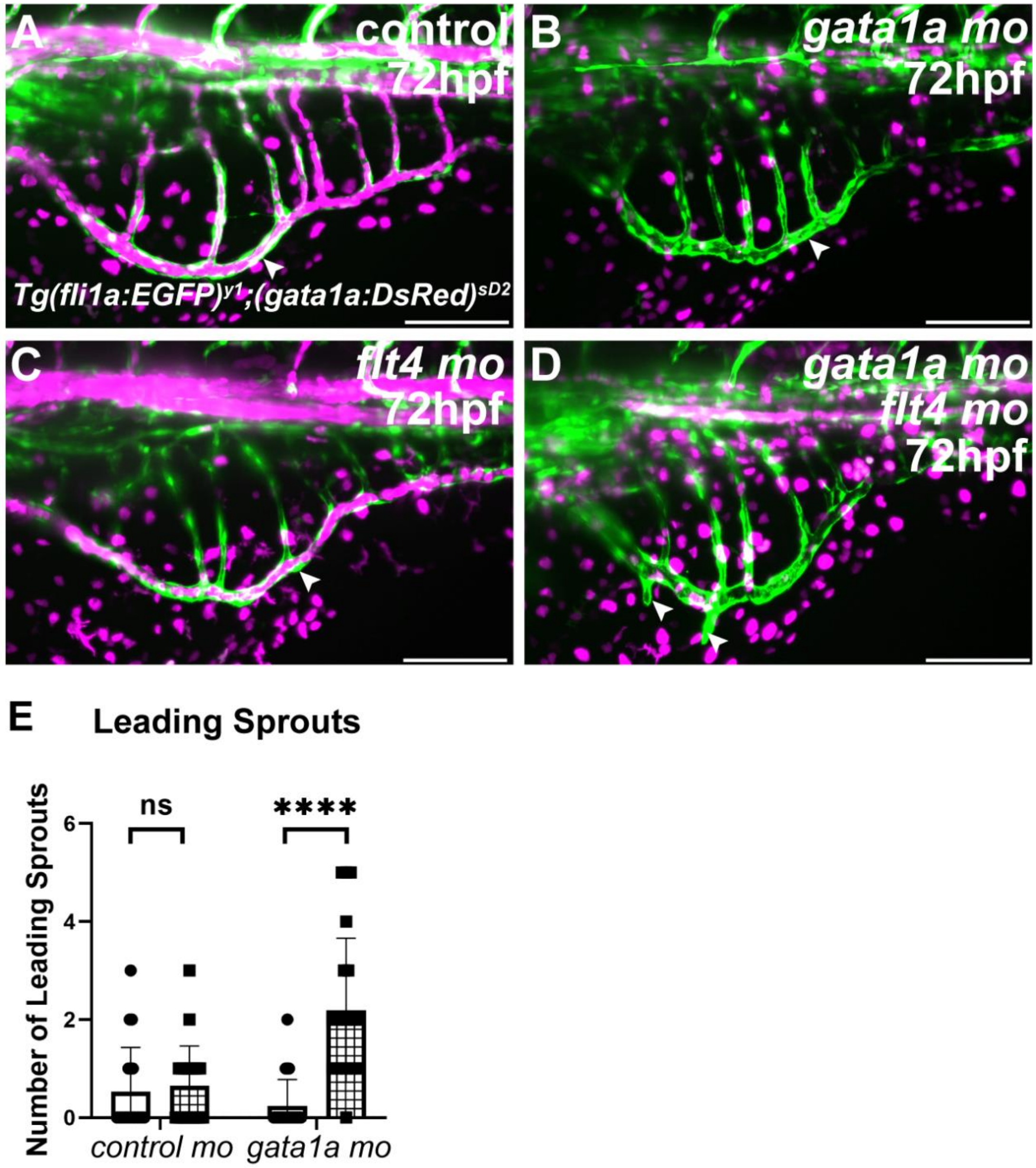


**Fig. S7 Leading sprout regression under low blood flow conditions is dependent on *flt4***

In comparison to controls (**A**), *gata1a* morphants displayed reduced circulating erythrocytes within the SIVP without inhibiting leading sprout regression (**B**, arrows). Leading sprout regression was unaltered by *flt4* knockdown under normal flow conditions (**C**) but *flt4* knockdown inhibited sprout regression under low flow conditions (**D**, arrows). The frequency of leading sprouts was significantly increased in *flt4*/*gata1a* double morphants compared to either *flt4*, *gata1a* or control single morphants (**E**). Two-way ANOVA, **** $p \leqslant 0.0001$; *** $p \leqslant 0.001$; ** $p \leqslant 0.01$; ns $p \geq 0.05$. control morphants n=19; *flt4* morphants n=19; *gata1a* morphants n=21; *flt4; gata1a* morphants n=21, 3 replicates. Scale bars 100μm.

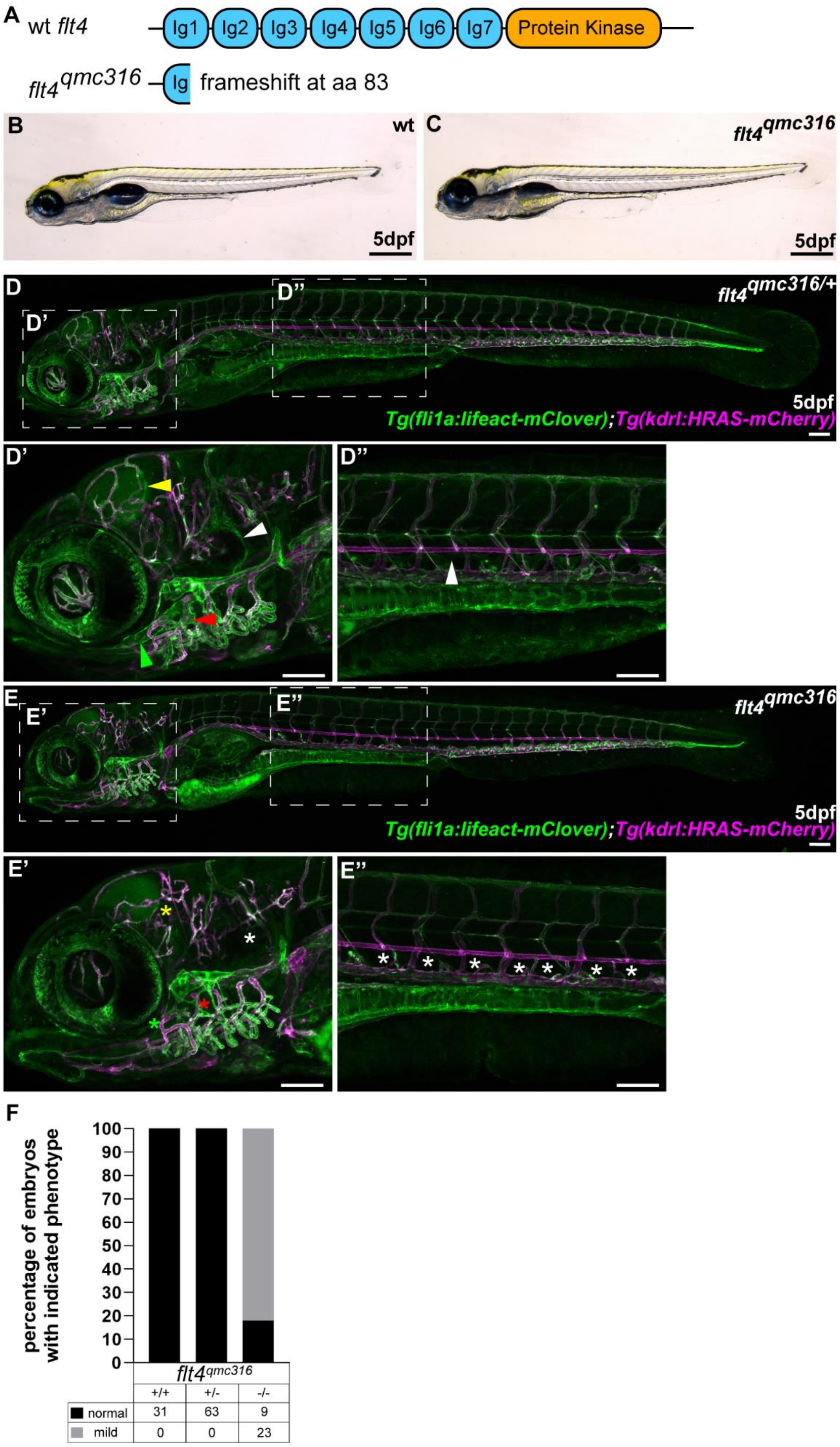

A
wt flt4
Ig1
Ig2
Ig3
Ig4
Ig5
Ig6
Ig7
Protein Kinase
flt4qmc316
Ig
frameshift at aa 83
B
wt
5dpf
C
flt4qmc316
5dpf
D
D'
D''
flt4qmc316/+
5dpf
Tg(fli1a:lifeact-mClover);Tg(kdrl:HRAS-mCherry)
D'
D''
E
E'
E''
flt4qmc316
5dpf
Tg(fli1a:lifeact-mClover);Tg(kdrl:HRAS-mCherry)
E'
E''
F
percentage of embryos with indicated phenotype
100
90
80
70
60
50
40
30
20
10
0
flt4qmc316
+/+
+/-
-/-
normal
31
63
9
mild
0
0
23

**Fig. S8 *flt4$^{qmc316}$* mutants display impaired lymphatic development.**

The *flt4$^{qmc316}$* mutant allele causes a premature truncation in the first immunoglobulin domain of Flt4, caused by a frameshift at amino acid 83 (**A**) *flt4$^{qmc316}$* mutants display no embryonic oedema by 5dpf compared to wild type siblings (**B-C**). *flt4$^{qmc316/+}$* heterozygotes display stereotypical formation of embryonic lymphatic vasculature (**D**), including meningeal lymphatic endothelial cells (**D'** yellow arrowhead), the otolithic facial lymphatic vessel (**D'** white arrowhead), the medial facial lymphatic vessel (**D'** red arrowhead) and the lateral facial lymphatic vessel (**D'** green arrowhead), and formation of the thoracic duct (**D''** white arrowhead). Conversely, *flt4$^{qmc316}$* mutants display impaired lymphatic vasculature formation (**E**), including loss of facial lymphatic vasculature (**E'** asterisks) and failure to form the thoracic duct (**E''** asterisks). Compared to wild type and heterozygous siblings (100%), 90% of *flt4$^{qmc316}$* mutants were identifiable at 5 dpf by a slight thickening of the membrane behind the eye (**F**). Scale bars in (B, C) = 500μm, scale bars in (D-E'') = 100μm
